# mGluR4–Npdc1 complex mediates α-synuclein fibril-induced neurodegeneration

**DOI:** 10.1101/2025.07.16.665194

**Authors:** Azucena Perez-Canamas, Mingming Chen, Leire Almandoz-Gil, Nabab Khan, Si Jie Tang, Allyson Ho, Erik C. Gunther, Stephen M. Strittmatter

**Author notes:** These authors contributed equally (Azucena Perez-Canamas, Mingming Chen).

## Abstract

Fibrils of misfolded α-synuclein (α-syn) accumulate in Parkinson’s disease and other synucleinopathies, spreading between cells to template further misfolding and drive neurodegeneration. α-syn fibril entry into healthy neurons is recognized as a key step in the disease process but remains ill-defined mechanistically. Here, we comprehensively assessed the membrane proteome for binding of α-syn fibrils. Expression cloning identified mGluR4 and Npdc1 as plasma membrane proteins expressed by substantia nigra neurons capable of supporting high affinity α-syn fibril binding. Moreover, mGluR4 and Npdc1 cellular signaling functions were titrated by the presence of extracellular fibrillary α-syn. While striatal α-syn fibril injection led to nigral dopamine neuron loss in wild type mice, deletion of either *Grm4* or *Npdc1* provided protection of dopamine neurons. We observed mGluR4 and Npdc1 to form a complex that regulates mGluR4 signaling. Cultured neurons lacking both *Grm4* and *Npdc1* fail to bind α-syn fibrils, to accumulate phosphorylated α-syn and to lose synapses. Transheterozygous *Grm4*, *Npdc1* mice showed protection from nigral neuron loss after striatal α-syn injection, demonstrating genetic interaction between the two binding proteins. On a transgenic α-syn A53T background, double *Grm4*, *Npdc1* heterozygosity robustly increased mouse survival, motor function and spinal motoneuron number. Thus, a cell surface mGluR4–Npdc1 complex participates in α-syn neurodegeneration.

---

Parkinson’s disease (PD), Diffuse Lewy Body disease (DLB) and related synucleinopathies are characterized by the accumulation of α-syn amyloid fibrils in Lewy bodies (LB) and Lewy neurites^1–3^. Multiple lines of evidence indicate that misfolded α-syn contributes to degenerative pathophysiology and symptoms^4^. Specifically, fibrillary α-syn amyloids can serve as templates to induce native α-syn monomers to misfold and propagate aberrant protein conformation^5–8^. Importantly, propagation can spread from the intracellular space of one cell to the intracellular space of another cell. Several models for cell spreading of α-syn and aggregates have been proposed, including tunneling nanotubes and bulk micropinocytosis or endocytosis^9, 10^. The most direct and selective model involves high-affinity receptor-mediated uptake of low concentrations of α-syn aggregates from the extracellular space into recipient cells for α-syn templating and neuronal dysfunction. The well documented ability of focal α-syn fibril injection into brain to spread misfolding and toxicity along neuronal pathways supports the latter hypothesis^5, 7, 8, 11^. However, the molecular basis for α-syn amyloid interaction with the neuronal cell surface to trigger aberrant neuronal signaling, internalization, and propagation remains poorly defined.

Previously, we utilized unbiased genome-wide receptor expression cloning to identify cellular prion protein (PrP^C^) as a high affinity receptor for misfolded forms of amyloid-ß in Alzheimer’s disease^12^. A parallel approach limited to 352 cDNAs was applied to identify binding sites for α-syn preformed fibrils (PFF) ^13^, the toxic aggregates prepared from monomeric α-syn protein. This revealed an affinity of α-syn PFF for LAG3^14^, although subsequent studies have indicated that LAG3 is expressed primarily by innate immune cells in the brain and may not be required for α-syn pathology in neurons^15^. Other potential α-syn fibril receptors have been proposed without unbiased screening. Prion protein itself was tested based on an analogy of misfolded α-syn with misfolded amyloid-ß^16^, but has yielded conflicting results^17, 18^. The PD risk gene, *GPNMB*, was postulated to encode an α-syn PFF binding site^19^, though its primary expression and action may be glial^20^, similar to LAG3. The microglial Fc-gamma receptor IIB (FcgRIIB) has also been implicated in extracellular α-syn toxicity and the spread of templated misfolding^21, 22^.

Here, we pursued a comprehensive screen for human membrane-associated proteins capable of supporting high affinity α-syn PFF binding to heterologous cells. A small number of proteins were confirmed as binding sites, and two were prioritized based on neuronal expression in dopamine cells (DA) of the substantia nigra, pars compacta (SNc), namely metabotropic glutamate receptor 4 (mGluR4) and Neuronal Proliferation, Differentiation and Control 1 (NPDC1). We show that these proteins bind α-syn PFF, that PFF binding alters their receptor function, and that they are essential for α-syn induced neurodegeneration. Unexpectedly, mGluR4 and NPDC1 physically and genetically interact in a complex which has prominent role in mediating α-syn PFF triggered neurodegeneration.

## Screen for α-syn fibril binding sites identifies nigral membrane proteins mGluR4 and NPDC1

Different membrane proteins have been proposed to act as receptors for α-syn aggregates^23^ but a comprehensive screen containing the majority of membrane proteins has not been reported. Here, we combined three different cDNAs expression libraries encoding more than 80% of all proteins classified by Gene Ontology with the term “plasma membrane” but excluding olfactory receptors (Extended Data File 1). Expression vectors for a total of 4,401 genes were individually transfected into HEK293T cells. Two days after transfection, binding assays were performed using biotinylated α-syn preformed fibrils (PFF) (Figure 1a; Extended Data Figure 1). As a result, we identified 16 novel membrane proteins that bind α-syn PFF (Extended Data Figure 2). From this list, we deprioritized B3GAT1 and CHST2 because they function primarily as intracellular synthetic enzymes in the secretory pathway. We also deprioritized CX3CL1 because this extracellular ligand is a peripheral membrane protein without a transmembrane segment. The remaining hit genes as well as several literature α-syn PFF binding proteins were assessed for expression in mouse midbrain DA cells using the https://brain-map.org website. Most of these genes were not expressed in nigral neurons, but *Grm4*, *Npdc1*, *Prnp* and *Lrrc4c* expression signals were clear (Extended Data Figure 3). A study of human nigral gene expression confirmed *GRM4*, *NPDC1* and *PRNP* expression, but not *LRRC4C* expression^24^. Since previous studies have disputed a role for PrP^C^ in synucleinopathy, we focused on mGluR4 and NPDC1 here. Of note, mGluR4 has also been considered a target for symptomatic treatment of motor symptoms in PD^25, 26^.

**Figure 1.**
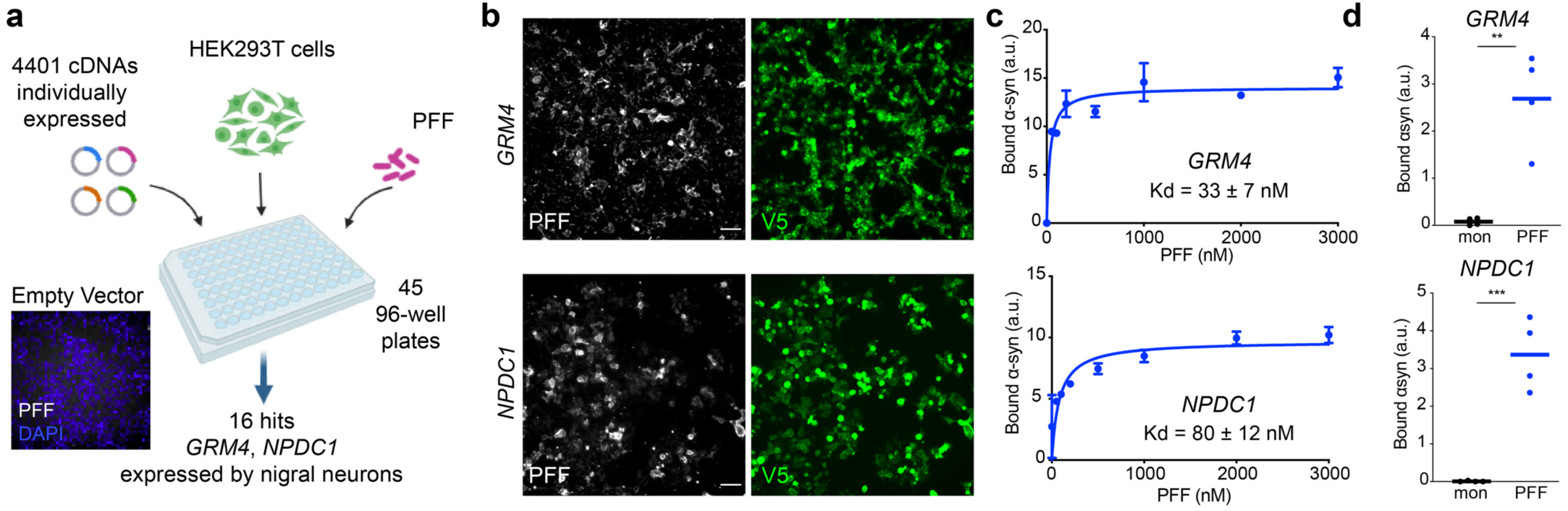
Comprehensive unbiased screen reveals GRM4 and NPDC1 as novel PFF-interactors. **a**, Schematic representation of the screening process. Briefly, HEK293T were plated in 96-well plates. A total of 4,401 cDNAs were used individually to transfect the cells. Two days after transfection, cells were treated with biotinylated α-syn PFF (1000 nM, monomer equivalents) for 2 h at RT and bound α-syn assessed. The micrograph shows that there was no detectable binding to control HEK293T cells. As a result, 16 membrane proteins were confirmed as PFF interactors. **b**, Representative images of HEK293T cells transfected with *GRM4* (upper panels) and *NPDC1* (lower panels) treated with biotinylated α-syn PFF (1000 nM) for 2 h and stained for α-syn PFF binding (streptavidin, gray) and GRM4 or NPDC1 expression (V5 tag, green). Scale bar = 100 µm. **c**, Graphs show mean ± SEM of the mean intensity of streptavidin (PFF) at different concentrations of α-syn PFF in HEK293T cells transfected with *GRM4* or *NPDC1*. N = 3 separate determinations. **d,** Graphs show mean ± SEM of the mean intensity of streptavidin (PFF) bound to HEK293T cells transfected with *GRM4* (upper graph) and *NPDC1* (lower graph) after exposure to 3000 nM α-syn monomer or PFF for cell transfected with *GRM4* (upper graph) and *NPDC1* (lower graph). **p<0.01, ***p<0.001, two tailed *t*-test.

Binding of α-syn PFF to mGluR4 and NPDC1 in HEK293T cells was saturable, with apparent disassociation constants of 33 nM and 80 nM, respectively, expressed as monomer equivalents (Figure 1b,c). With fibrils sonicated to a length of 50 nm, the concentration of α-syn PFF is at least 100-fold lower that the monomer equivalent, in the picomolar range. Binding was specific for the misfolded α-syn conformation, since cells expressing these proteins did not bind α-syn monomers to an extent greater than background at concentrations up to 3 µM (Fig. 1d). Binding to mGluR4 was specific amongst mGluRs, since mGluR1, 2, 3, 5 and 8, were negative for α-syn PFF interaction (A.P.C. and S.M.S., unpublished observations). Co-immunoprecipitation analysis in HEK293T cells overexpressing mGluR4 and NPDC1 revealed a direct physical interaction of both proteins with α-syn PFF (Extended Data Figure 4). Thus, a comprehensive unbiased screen of membrane proteins identified 16 previously unrecognized proteins that interact with α-syn PFF. mGluR4 and NPDC1 were selected as candidates for further analysis as potential mediators of α-syn PFF toxicity in nigral dopamine neurons.

## α-syn fibrils regulate mGluR4 and NPDC1 cellular functions

Next, we checked whether α-syn PFF binding to mGluR4 and NPDC1 altered physiological receptor function. mGluR4 is a metabotropic glutamate receptor that signals through the inhibition of cyclic AMP (cAMP) cascade after activation with glutamate ligand^27^. To analyze the effect of α-syn PFF on mGluR4 function, we performed a calcium assay in HEK293T cells stably expressing mGluR4 and Gαqi5, a chimeric Gq subunit that allows stimulation of phospholipase C by receptors otherwise coupled exclusively to Gi^28^. Intracellular calcium measurements using L-AP4 as an agonist revealed an increase in calcium response when cells were treated with a low dose of α-syn PFF (62.5 nM monomer equivalent) compared to controls (DPBS and monomeric α-syn) (Figure 2a). However, when using a higher α-syn PFF dose (1,000 nM monomer equivalent) the outcome was opposite: α-syn PFF blocked calcium signaling. Dose titration indicated that α-syn PFF concentrations above 125 nM blocked subsequent mGluR4 activation by L-AP4 (Figure 2b).

**Figure 2.**
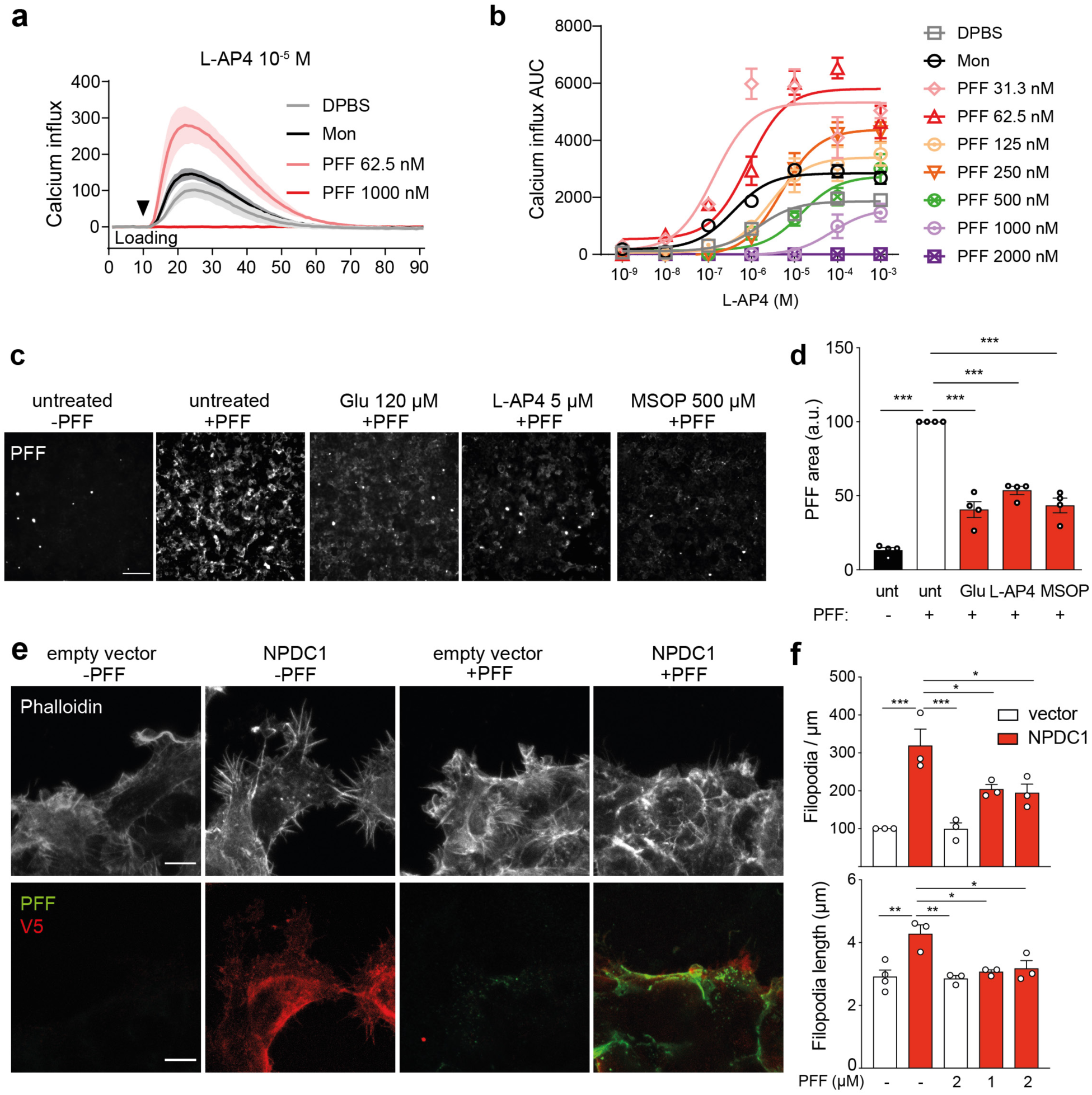
α-syn PFF interaction with GRM4 and NPDC1 impairs receptor function. **a,** Graph shows the calcium influx as a response to L-AP4 addition (10^-5^ M) in stable GRM4-Gαqi5 cells treated with DPBS, α-syn monomer or α-syn PFF (62.5 nM and 1000 nM). Mean ± SEM indicated from 9 separate wells. **b,** Graph shows mean ± SEM of the area under the curve in a calcium assay response to different concentrations of L-AP4 in stable GRM4-Gαqi5 cells treated with DPBS, α-syn monomer or PFF at different concentrations. N=3 separate experiments. ***p<0.001. **c,** Representative images of HEK293T cells transfected with *GRM4* and treated with mGluR4 ligands glutamate, L-AP4 and MSOP at the indicated concentrations before the addition of biotinylated α-syn PFF for 3 h. α-syn PFF bound to the cells are detected with streptavidin. Scale bar = 200 µm. **d,** Graph shows mean ± SEM of the area occupied with PFF in HEK293T treated with biotinylated α-syn PFF for 3 h after the addition of mGluR4 ligands (Glu, L-AP4 and MSOP). One-way ANOVA with Tukey’s post-hoc test. **e,** Representative images of HEK293T cells transfected or not with *NPDC1* and treated with biotinylated α-syn PFF for 3 h. Cells were stained with phalloidin (grey, to detect actin) and V5 (green, to confirm NPDC1 expression). α-syn PFF bound to the cells are detected with streptavidin (red). Scale bar = 10 µm. **f,** Graph shows mean ± SEM of the number of filopodia per µm (top) and filopodia length (bottom) in HEK293T cells treated with biotinylated α-syn PFF for 3 h. One-way ANOVA with Tukey’s post hoc test. *p<0.05, **p<0.01, ***p<0.001.

We also checked whether the presence of mGluR4 orthosteric agonists (glutamate and L-AP4) or the orthosteric antagonist MSOP affect α-syn PFF binding to the receptor. Binding assays in HEK293T cells overexpressing mGluR4 and treated with the different orthosteric ligands showed a decrease in α-syn PFF binding compared to untreated controls (Figure 2c,d), indicating antagonism between known mGluR4 ligands and α-syn PFF binding. The reduction in α-syn PFF binding was not due to alterations in the plasma membrane levels of mGluR4 upon ligand addition (Extended Data Figure 5a,b) nor to changes in total amount of mGluR4 present (Extended Data Figure 5c,d) even when using high concentrations of drugs (Extended Data Figure 5e,f). Treatment with the same compounds in NPDC1-overexpressing cells showed no effect in α-syn PFF binding or total NPDC1 expression (Extended Data Figure 5g-i), consistent with a receptor-specific effect on mGluR4 conformational state.

NPDC1 was originally identified in a neuronal precursor cell line after growth arrest, indicating it may inhibit proliferation^29^. NPDC1 is a single transmembrane protein where the N-terminal (Nt) part is extracellular and C-terminal (Ct) intracellular^30^. We deleted two regions of the extracellular domain (Extended Data Figure 5j) in order to identify the interacting region with α-syn PFF. Both deletions abrogated binding of α-syn PFF (Extended Data Figure 5k,l) without major changes in NPDC1 expression (Extended Data Figure 5m).

When performing binding assays in HEK293T cells, we noted that NPDC1-overexpressing cells displayed increased numbers of filopodia, and that these were enriched for filamentous actin detected with rhodamine-phalloidin (Figure 2e,f). Thus, we assessed whether α-syn PFF binding altered this phenotype induced by NPDC1 expression. Addition of α-syn PFF to NPDC1-overexpressing cells caused a decrease in the number and length of filopodia compared to non-treated cells (Figure 2e,f). Thus, expression of NPDC1 induces an increase in number and length of filopodia, which is prevented in the presence of α-syn PFF as an NPDC1 ligand.

While these studies demonstrate α-syn PFF regulation of mGluR4 and NPDC1, activity of the deprioritized candidate LRRC4C was not altered by α-syn PFF. Specifically, binding of a soluble form of the LRRC4C ligand netrin-G1 was not altered by PFF, and the synaptogenic activity of LRRC4C expressed in HEK cells for cortical neurons^31^ was not altered by the presence of α-syn PFF (L.A.-G. and S.M.S., unpublished observations). Based on these interaction studies, we explored the potential role of mGluR4 and NPDC1 in α-syn mediated neurodegeneration in mice.

## *Grm4* is required for nigral dopamine neuron loss induced by α-syn fibril striatal injection

In order to evaluate the role of mGluR4 in α-syn induced dopamine neurodegeneration *in vivo*, we performed intrastriatal injections of α-syn PFF^5^ in wt and *Grm4*-/- mice^32^ and analyzed histology and behavior 6 months after the injections (Figure 3a). General mouse health was not altered as body weights were unchanged in both genotypes after α-syn PFF injections (Extended Data Figure 6a). In the striatum, the injection resulted in local α-syn phosphorylation (p-α-syn) and aggregation in both strains (Figure 3b,d; Extended Data Figure 6c). The staining of tyrosine hydroxylase (TH) immunoreactive fibers in the striatum was not significantly reduced in the wt mice injected with α-syn PFF as compared to PBS, and was similar in *Grm4*-/- mice (Figure 3b,c; Extended Data Figure 6b).

**Figure 3.**
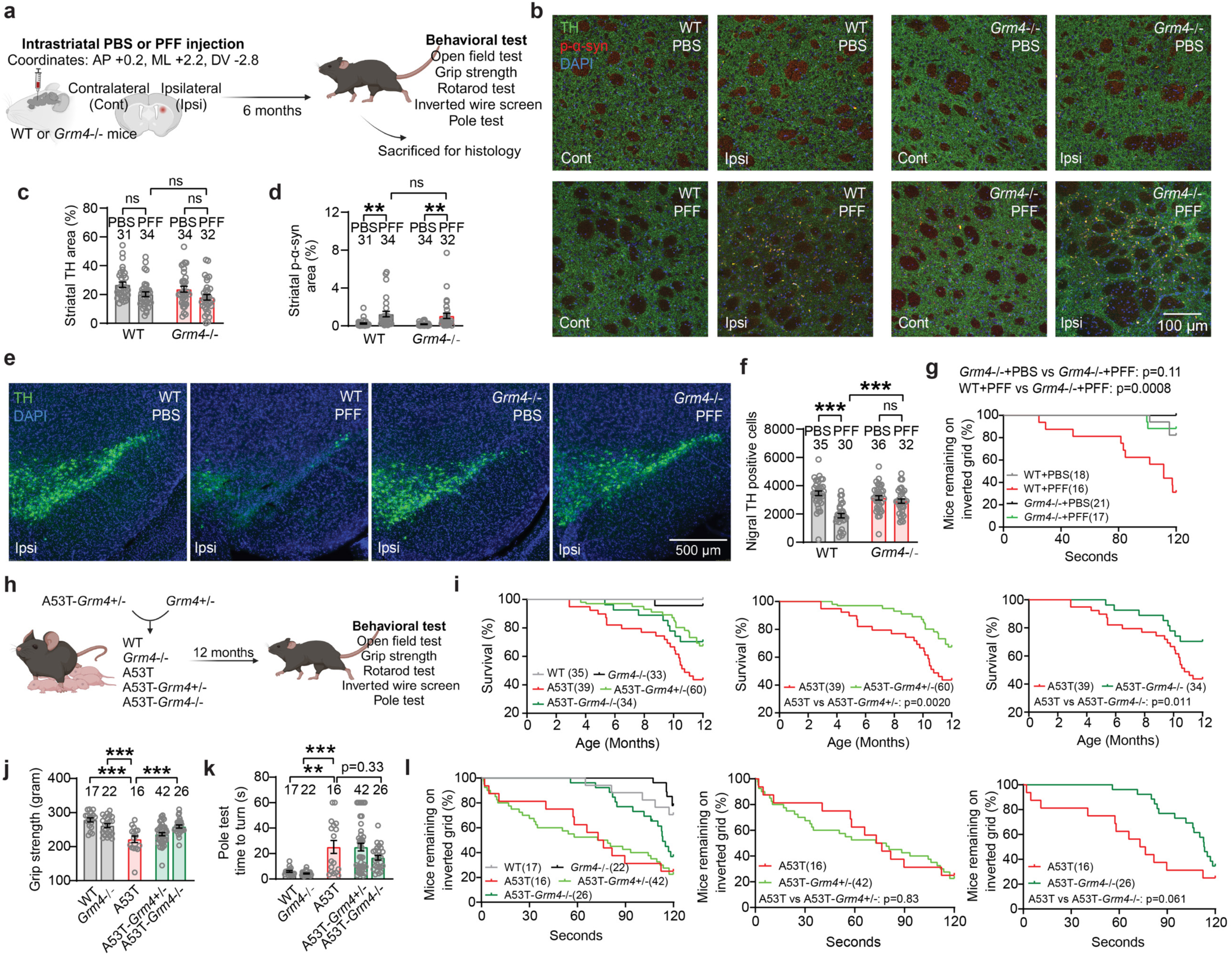
*Grm4* deletion protects nigral neurons and motor function after striatal α-syn PFF injection and improves α-syn A53T mouse function. **a,** Diagram illustrating the experimental design for α-syn PFF injection into wt and *Grm4*-/- mice striatum. **b,** Representative images showing TH and p-α-syn co-staining in the caudate-putamen of wt and *Grm4*-/- mice injected with α-syn PFF. Scale bar = 100 µm. **c,** Graph shows mean ± SEM of TH levels in the striatum of wt and *Grm4*-/- mice injected with α-syn PFF. One-way ANOVA with by Tukey’s post hoc test. **d,** Graph shows mean ± SEM of p-α-syn levels in the striatum of wt and *Grm4*-/- mice injected with α-syn PFF. One-way ANOVA with by Tukey’s post hoc test. **p=0.0052 for WT-PBS vs WT-PFF; **p=0.0098 for Grm4 KO-PBS vs Grm4 KO-PFF. **e,** Representative images of TH staining in the ipsilateral side of the SNc of wt and *Grm4*-/- mice injected with α-syn PFF. Coronal section with dorsal up and midline to left. Scale bar = 500 µm. **f,** Graph shows mean ± SEM of TH positive neurons in the ipsilateral side of the SNc of wt and *Grm4*-/- mice injected with α-syn PFF. One-way ANOVA with by Tukey’s post hoc test. ***p<0.0001 for WT-PBS vs WT-PFF and WT-PFF vs Grm4 KO-PFF, mean ±SEM. **g,** Graph shows the percentage of wt and *Grm4*-/- mice injected with α-syn PFF remaining in the inverted grid in the wire screen test. Gehan-Breslow-Wilcoxon test. **h,** Diagram illustrating the experimental design for generating and analyzing A53T-*Grm4*-/- mice. **i,** Graphs show survival rates for A53T-*Grm4*-/- mice. Gehan-Breslow-Wilcoxon test. **j,** Graph shows mean ± SEM of grip strength in A53T-*Grm4*-/- mice. One-way ANOVA with Tukey’s post hoc test. ***p<0.0001 for WT vs A53T, Grm4 KO vs A53T, and A53T vs A53T-Grm4 KO. **k,** Graph shows mean ± SEM of time to turn in the pole test in A53T-*Grm4*-/- mice. One-way ANOVA with Tukey’s post hoc test. ***p<0.0001 for Grm4 KO vs A53T; **p=0.0015 for WT vs A53T. **l,** Graphs show the percentage of A53T-*Grm4*-/- mice remaining in the inverted grid in the inverted wire screen test. Gehan-Breslow-Wilcoxon test.

Stereological counts of dopaminergic TH+ cells in the SNc revealed a robust decrease in dopaminergic neurons in the ipsilateral site of wt mice injected with α-syn PFF (Figure 3e,f; Extended Data Figure 6d). Dopaminergic cell loss was prevented in *Grm4*-/- injected mice, indicating that mGluR4 is required for cell loss induced by α-syn PFF. Significant neuronal loss was not observed in the contralateral SNc for any cohort (Extended Data Figure 6d-f).

We also analyzed the effect of mGluR4 knockout on behavioral consequences of α-syn PFF injection. The wire screen test measures motor abilities using an inverted grid. In this test, α-syn PFF injection impaired performance of wt mice, but not of *Grm4*-/- mice (Figure 3g), indicating that mGluR4 is required for this α-syn PFF-induced motor impairment. Other tests performed, including vertical pole test, open field, measurement of grip strength and rotarod, did not show a significant effect of α-syn PFF administration on motor behavior in wt or *Grm4*-/- mice (Extended Data Figure 6g-m).

To confirm and extend these results we employed a genetic model, the A53T transgenic mouse^33^, which expresses a A53T missense mutant form of human α-syn under the control of the murine prion promoter. A53T mice present an age-dependent phenotype including progressive motor deficits, intraneuronal inclusion bodies and neuronal loss most prominent in the spinal cord^33, 34^. Thus, we crossbred A53T transgenic mice with *Grm4*-/- mice in order to obtain double mutants and conducted behavioral tests at 12 months of age (Figure 3h). A53T mice showed a slight decrease in body weight when compared to wt littermate controls (Extended Data Figure 6n). Survival analysis of the different genotypes revealed a significant decrease in survival for A53T mice compared to wt littermate controls (Figure 3i), as reported^33, 34^. Survival of A53T-Grm4 KO was greater than that for A53T mice (Figure 3i), indicating that mGluR4 participates in the early death of A53T mice.

At 12 months of age, surviving A53T mice showed a reduction in grip strength compared to wt and *Grm4*-/- mice. The A53T grip strength decrement was alleviated in surviving A53T-Grm4 KO, but not in surviving A53T-Grm4 HET (Figure 3j). A deficit in motor abilities was also observed in A53T mice when performing the vertical pole test. This A53T motor deficit relative to wt mice was not corrected in A53T-Grm4 HET and A53T-Grm4 KO mice (Figure 3k). In the wire screen test, the percentage of surviving A53T and A53T-Grm4 HET mice remaining on the grid as a function of time was reduced compared to the other genotypes (Figure 3l), indicating a motor impairment for these groups. The A53T-Grm4 KO mice remained on the grid for a numerically longer time than A53T mice (p = 0.061). In the open field test, A53T-Grm4 KO mice were slightly faster than the other groups (Extended Data Figure 6o,p), and no differences were observed among the different genotypes in the rotarod test (Extended Data Figure 6q). Altogether, these results suggest that mGluR4 deletion reduces a subset of motor abnormalities present on the A53T background as well as protecting nigral DA neurons from α-syn PFF induced degeneration.

## Mice lacking *Npdc1* protect nigral neurons from α-syn fibril pathology

We used a parallel experimental design to investigate potential Npdc1 effects in α-syn pathology *in vivo* after α-syn PFF injection. Mice were assessed 6 months after the injections (Figure 4a). Body weight monitoring showed that wt and *Npdc1*-/- mice ^35^ remained healthy after α-syn PFF injections (Extended Data Figure 7a). Similar to the *Grm4*-/- cohort, α-syn PFF delivery did not decrease striatal TH levels in wt mice injected with α-syn PFF (Figure 4b,c; Extended Data Figure 7b). The local striatal increase in α-syn phosphorylation after α-syn PFF injections was similar in wt and *Npdc1*-/- mice (Figure 4b,d; Extended Data Figure 7c).

**Figure 4.**
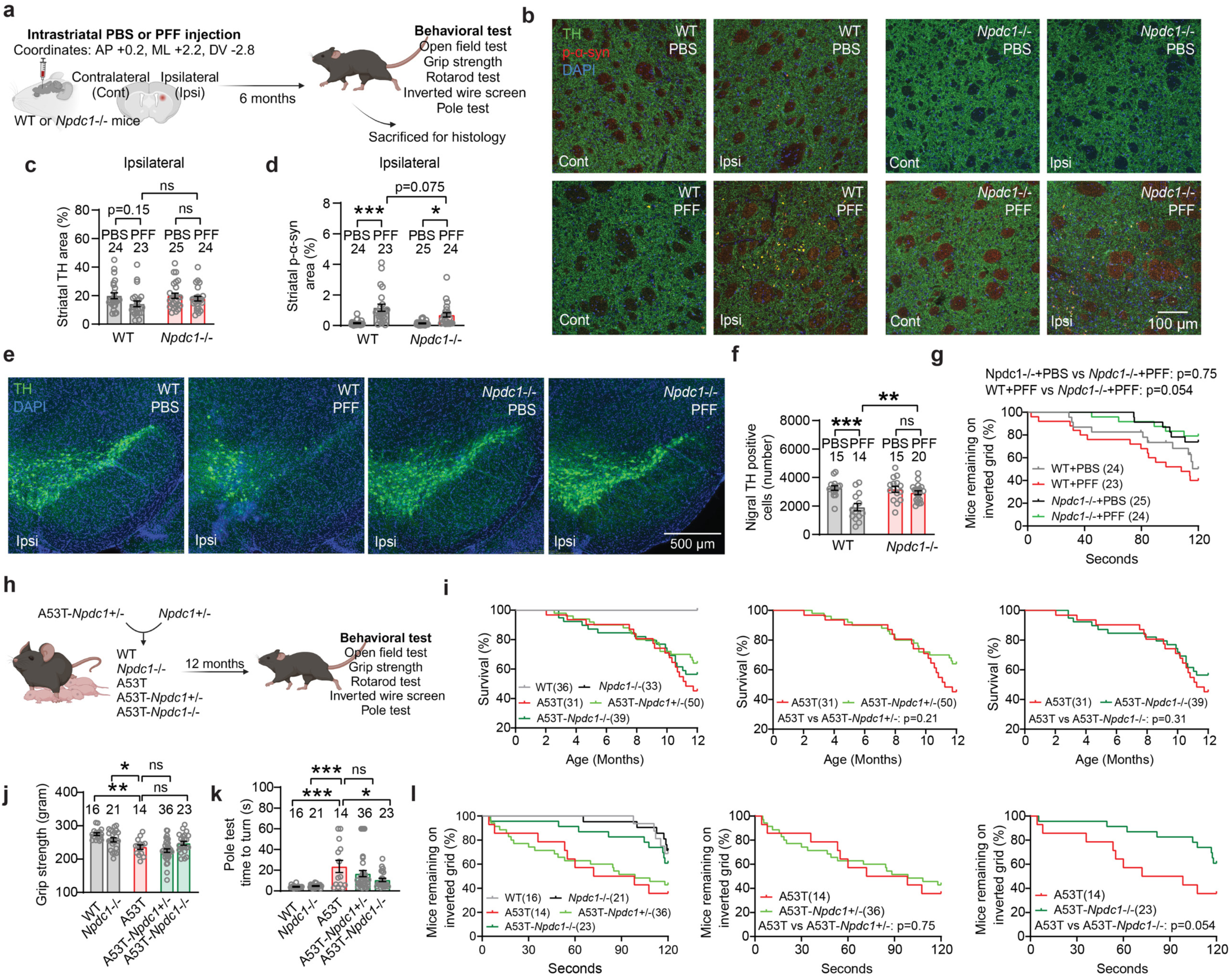
*Npdc1* deficiency limits nigral cell loss and improves motor function after striatal α-syn PFF injection and α-syn A53T expression. **a,** Diagram illustrating the experimental design for α-syn PFF injection into wt and *Npdc1*-/- mice striatum. **b,** Representative images showing TH and p-α-syn co-staining in the caudate-putamen of wt and *Npdc1*-/- mice injected with α-syn PFF. Scale bar = 100 µm. **c,** Graph shows mean ± SEM of striatal TH levels in the ipsilateral side of the striatum in wt and *Npdc1*-/- mice injected with α-syn PFF. One-way ANOVA with Tukey’s post hoc test. **d,** Graph shows mean ± SEM of striatal p-α-syn levels in the ipsilateral side of the striatum in wt and *Npdc1*-/- mice injected with α-syn PFF. One-way ANOVA with Tukey’s post hoc test. ***p<0.0001 for WT-PBS vs WT-PFF; *p=0.020 for Npdc1 KO-PBS vs Npdc1 KO-PPF. **e,** Representative images of TH staining in ipsilateral side of the SNc of wt and *Npdc1*-/- mice injected with α-syn PFF. Coronal section with dorsal up and midline to left. Scale bar = 500 µm. **f,** Graph shows mean ± SEM of TH positive neurons in the ipsilateral side of the SNc of wt and *Npdc1*-/- mice injected with α-syn PFF. One-way ANOVA with Tukey’s post hoc test. ***p<0.0001 for WT-PBS vs WT-PFF; **p=0.0014 for WT-PFF vs Npdc1 KO-PPF. **g,** Graph shows the percentage of wt and *Npdc1*-/- mice injected with α-syn PFF remaining in the inverted grid in the wire screen test. Gehan-Breslow-Wilcoxon test. **h,** Diagram illustrating the experimental design for generating and analyzing A53T-*Npdc1*-/- mice. **i,** Graphs show survival rates for A53T-*Npdc1*-/- mice. Gehan-Breslow-Wilcoxon test. **j,** Graph shows mean ± SEM of grip strength in A53T-*Npdc1*-/- mice. One-way ANOVA with Tukey’s post hoc test. **p=0.0023 for WT vs A53T; *p=0.030 for Npdc1 KO vs A53T. **k,** Graph shows mean ± SEM of time to turn in the pole test in A53T-*Npdc1*-/- mice. One-way ANOVA with Tukey’s post hoc test. ***p<0.0001 for WT vs A53T and Npdc1 KO vs A53T; *p=0.034 for A53T vs A53T-Npdc1 KO. **l,** Graphs show the percentage of A53T-*Npdc1*-/- mice remaining in the inverted grid in the wire screen test. Gehan-Breslow-Wilcoxon test.

Critically, α-syn PFF injection resulted in dopaminergic neuronal loss from the ipsilateral SNc of these wt mice, but not of *Npdc1*-/- mice (Figure 4e,f; Extended Data Figure 7d). Thus, Npdc1 depletion is protective against the nigral DA cell loss induced by α-syn PFF *in vivo*. No neuronal loss was observed in the contralateral site for any of the genotypes (Extended Data Figure 7d-f). In the wire screen test, PFF-injected *Npdc1-/-* remained on the grid for an intermediate period, not significantly different from PBS-injected *Npdc1-/-* mice or from PFF-injected wt mice (Figure 4g). α-syn PFF delivery had no effect in the vertical pole test, open field, measurement of grip strength and rotarod in the control wt mice, or in the *Npdc1*-/- mice (Extended Data Figure 7g-k).

*Npdc1*-/- mice were also crossbred with A53T transgenic in order to obtain double mutants. As for the A53T-Grm4 cohort, behavioral tests were conducted at 12 months of age (Figure 4h). In this cohort, A53T mice showed a trend to decrease in body weight, not significantly different from wt littermate controls (Extended Data Figure 7l). Survival analysis of the different genotypes revealed a significant decrease in survival rate of A53T mice compared to control, and this was not improved by *Npdc1* deletion (Figure 4i).

Behaviorally, surviving A53T mice at 12 months of age showed a reduction in grip strength compared to wt and *Npdc1*-/- mice, and this was not lessened by *Npdc1* deletion (Figure 4j). In contrast, the impairment of A53T mice on the vertical pole test was rescued in A53T-Npdc1 KO mice (Figure 4k). In the wire screen test, the percentage of A53T and A53T-Npdc1 HET mice remaining on the grid was less than in the rest of the genotypes (Figure 4l), indicating a motor impairment in these two groups. The percentage of A53T-Npdc1 KO mice was greater than in A53T mice, though the difference was not statistically significant (p = 0.054). No differences were observed among the surviving mice of the different genotypes in the open field or rotarod tests (Extended Data Figure 6m-o). These data indicate that Npdc1 removal improves a subset of motor deficits present in A53T mice and also rescues nigral DA neurons from α-syn PFF induced degeneration.

## mGluR4 and NPDC1 form a complex regulating mGluR4 activation and mediating α-syn fibril action

While the protection of nigral DA neurons from α-syn PFF induced degeneration is robust with either mGluR4 or NPDC1 gene knockout, other phenotypes are only partially protected by single gene deletion. This suggested that the two proteins have partially redundant roles. We also noted that the benefits of *Npdc1* deletion on α-syn pathologies in mice are strikingly similar to the improvements seen in *Grm4* null mice. This suggested that the two proteins might function cooperatively, as a complex. To explore potential interaction between mGluR4 and NPDC1, we performed co-immunoprecipitation (co-IP) in HEK293T cells overexpressing both receptors. Retention of NPDC1 with V5 beads showed that NPDC1 co-immunoprecipitated with mGluR4, but not mGluR5, another metabotropic glutamate receptor, indicating that the mGluR4–NPDC1 interplay is specific (Figure 5a,b). Reverse co-IP experiments, retaining mGluR4 with DYDDK beads, confirmed an interaction of Npdc1 with mGluR4, but not mGluR5 (Figure 5a,b). No available antibody was adequate to assess association of endogenous brain proteins, so we evaluated functional interactions. We performed calcium assays in GRM4-Gαqi5 cells transfected or not with NPDC1. NPDC1 co-expression completely blocked the calcium response of mGluR4 cells to L-AP4 stimulation, indicating that the presence of NPDC1 negatively impacts mGluR4 function (Figure 5c, Extended Data Figure 8a,b). To examine whether NPDC1 interacts with mGluR4 function in *cis* and/or in *trans*, we performed the same calcium assay but with a mixture of GRM4-Gαqi5 cells with NPDC1-transfected cells. Mixing GRM4-Gαqi5 cells with HEK293T cells transfected with an empty vector did not alter calcium responses (Figure 5d, Extended Data Figure 8c,d). However, combining GRM4-Gαqi5 cells with NPDC1-expressing cells produced a significant reduction in calcium response after stimulation at a 1:1 ratio or a complete blockade of the response at a 1:2 ratio (Figure 5d, Extended Data Figure 8e,f). Thus, the expression of NPDC1 in adjacent cells is sufficient to block mGluR4 function in *trans*, although *cis* interactions may be equally effective.

**Figure 5.**
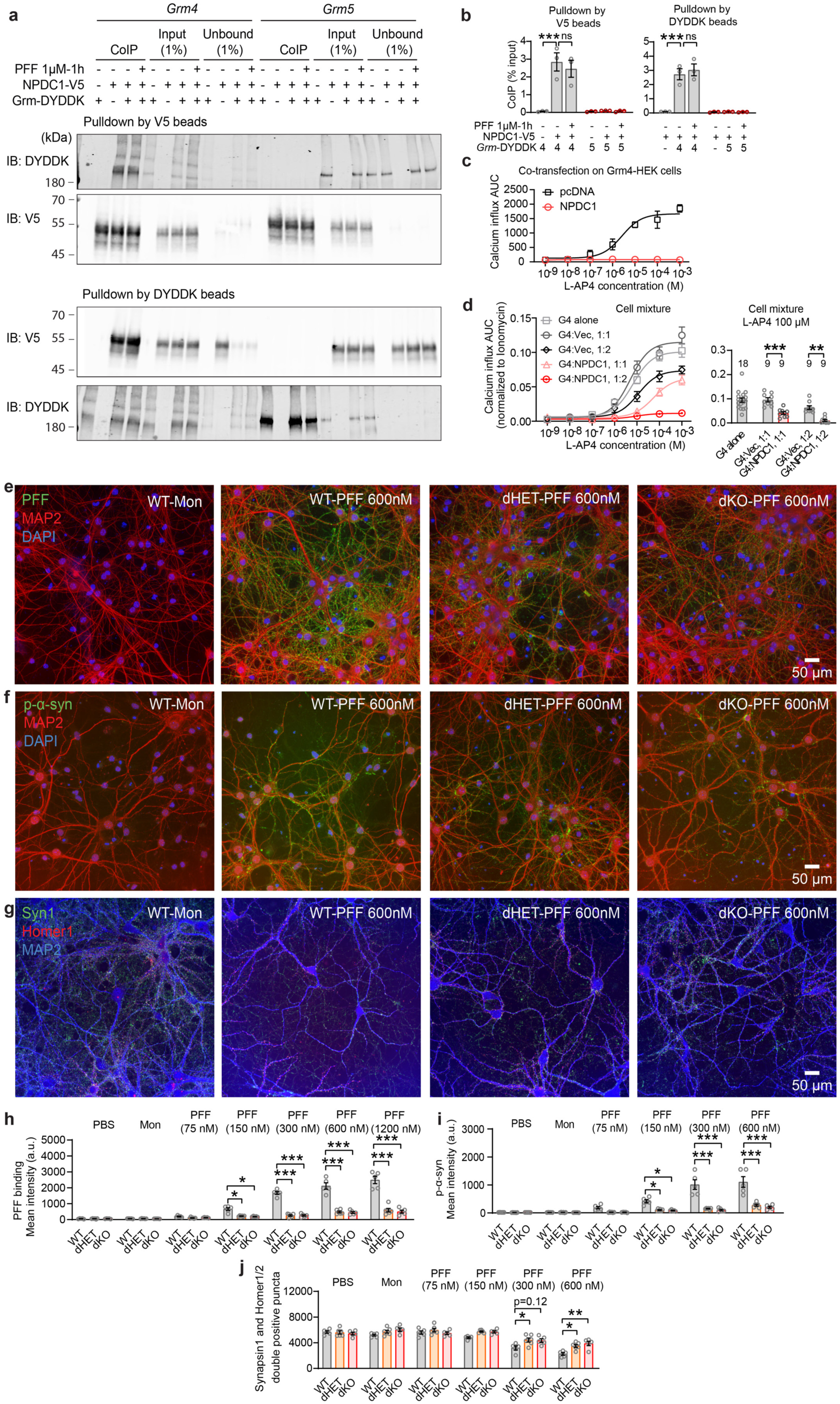
mGluR4 and Npdc1 physically associate and act synergistically to mediate PFF binding and synaptic loss in cortical neurons. **a,** Representative immunoblots with anti-V5 and anti-DYDDK of co-IP assays using HEK293T cells expressing Grm4-DYDDK or Grm5-DYDDK, together with V5-tagged NPDC1. **b,** Graphs show mean ± SEM of pulldown by V5 beads (left) and DYDDK beads (right). One-way ANOVA with Tukey’s post hoc test. For both V5 and DYDDK pulldown: ***p<0.0001 for Grm4-DYDDK vs NPDC1-V5 and Grm4-DYDDK. **c,** Graph shows mean ± SEM of calcium influx curve after L-AP4 stimulation in GRM4-Gαqi5 cells (G4) transfected with an empty vector (Vec, pcDNA) or with *NPDC1.* **d,** (left) Graph shows mean ± SEM of calcium influx curve after L-AP4 stimulation in GRM4-Gαqi5 cells (G4) co-cultured with HEK293T cells transfected with an empty vector (Vec, pcDNA) or with *NPDC1* at different ratios. **d**, (right) Graph shows mean ± SEM of calcium response after 100 µM L-AP4 stimulation in GRM4-Gαqi5 cells (G4) co-cultured with HEK293T cells transfected with an empty vector (Vec, pcDNA) or with *NPDC1* at different ratios. One-way ANOVA with Tukey’s post hoc test. ***p=0.0009 for G4:Vec, 1:1 vs G4:NPDC1, 1:1; **p=0.0015 for G4:Vec, 1:2 vs G4:NPDC1, 1:2; **e,** Representative images of streptavidin (PFF, green), MAP2 (red), and DAPI (blue) staining in cultured neurons from WT, dHET and dKO mice after 2 hours of exposure with biotinylated α-syn monomer or PFF at 37°C. Scale bar = 50 µm. **f,** Representative images of p-α-syn (green), MAP2 (red) and DAPI (blue) staining in cultured neurons from WT, dHET and dKO mice after 7 days of exposure with biotinylated α-syn monomer or PFF at 37°C. Scale bar = 50 µm. **g,** Representative images of Synapsin1 (green), Homer1/2 (red), and DAPI (blue) staining in cultured neurons from WT, dHET and dKO mice after 7 days of exposure with biotinylated α-syn monomer or PFF at 37°C. Scale bar = 50 µm. **h,** Graph shows mean ± SEM of streptavidin (PFF) mean intensity on MAP2 positive area in cultured neurons from WT, dHET and dKO mice after 7 days of exposure with biotinylated α-syn monomer or PFF at 37 °C. One-way ANOVA with Tukey’s post hoc test. For 150 nM: *p=0.018 for WT vs dHET, *p=0.012 for WT vs dKO. For 300 nM: ***p<0.0001 for WT vs dHET, ***p<0.0001 for WT vs dKO. For 600 nM: ***p<0.0001 for WT vs dHET, ***p<0.0001 for WT vs dKO. For 1200 nM: ***p<0.0001 for WT vs dHET, ***p<0.0001 for WT vs dKO. **i,** Graph shows mean ± SEM of p-α-syn mean intensity on MAP2 positive area in cultured neurons from WT, dHET and dKO mice after 7 days of exposure with biotinylated α-syn monomer or PFF at 37 °C. One-way ANOVA with Tukey’s post hoc test. For 150 nM: *p=0.034 for WT vs dHET, *p=0.028 for WT vs dKO. For 300 nM: ***p<0.0001 for WT vs dHET, ***p<0.0001 for WT vs dKO. For 600 nM: ***p<0.0001 for WT vs dHET, ***p<0.0001 for WT vs dKO. **j,** Graph shows mean ± SEM of Synapsin1 and Homer1/2 double positive dots on MAP2 positive area in cultured neurons from WT, dHET and dKO mice after 7 days of exposure with biotinylated α-syn monomer or PFF at 37°C. One-way ANOVA with Tukey’s post hoc test. For 300 nM: *p=0.040 for WT vs dHET. For 600 nM: *p=0.039 for WT vs dHET, **p=0.0014 for WT vs dKO.

Since mGluR4 and NPDC1 interact physically and functionally, we decided to investigate whether the elimination or reduction of both proteins synergized in neurons with regard to their effect on α-syn PFF binding, the accumulation of aggregated α-syn, and synaptic damage. We crossbred *Grm4*-/- and *Npdc1*-/- mice and cultured cortical neurons from wt, double heterozygous (dHET) and double ko (dKO) embryos. Incubation with α-syn PFF for 2 hours at 37 °C or 4 °C revealed a significant decrease in α-syn PFF binding in dHET and dKO compared to wt mice (Figure 5e,h, Extended Data Figure 8g,h) without alterations in total neuronal density as measured with MAP2 staining (Extended Data Figure 8g,i,j). The greater suppression of binding in the double knockout as compared single knockouts supports partially redundant action of these two α-syn PFF binding proteins. More strikingly, the strong reduction of binding to double heterozygous neurons documents a genetic interaction that supports a critical role for the mGluR4 – NPDC1 physical complex in neuronal binding of α-syn PFF.

When neurons are exposed to α-syn PFF for 7 days, aggregates of p-α-syn accumulate in these cells. Both dHET and dKO neurons showed a marked reduction in accumulation of p-α-syn (Figure 5f,i) without changes in neuronal density detected by MAP2 for PFF concentrations up to 600 nM (Extended Data Figure 8k,l). While neuronal cell density measured by MAP2 staining is not altered in this situation, exposure of wt neurons to α-syn PFF for 7 days led to reduced synaptic density as measured by colocalization of puncta stained for synapsin1 and Homer1/2. Neurons of both the dHET and dKO genotype were protected from synapse loss triggered by α-syn PFF (Figure 5g,j). In summary, mGluR4 and Npdc1 individually have partially redundant roles in α-syn PFF binding to neurons as evidenced by comparing dKO with single KO. Furthermore, the physical and genetic interaction of mGluR4 and NPDC1 is required for efficient binding, accumulation of pathological aggregates and neuronal toxicity as shown in the dHET phenotypes.

## Transheterozygote *Grm4, Npdc1* nigral neurons are protected from α-syn fibril striatal injection

Given the genetic interactions of mGluR4 and Npdc1 in culture, we investigated the impact of their combined absence *in vivo*. Littermate wt, dHET and dKO mice were injected with α-syn PFF in the striatum and analyzed 6 months after injection. No differences in body weight progression were observed among the groups (Extended Data Figure 10a). In line with the single knockout studies, α-syn PFF generated local accumulation of phosphorylated α-syn in the striatum but no change in striatal TH fibers from wt (Extended Data Figure 9b-d). The striatal effects of α-syn PFF were similar in all genotypes.

For the wt mice, the striatal α-syn PFF injection generated a pronounced decrease in the number of TH-positive (dopaminergic, DA) cells in the ipsilateral SNc (Figure 6a,b, Extended Data Figure 9d). Critically, removal of a single copy (dHET) or two copies (dKO) of both *Grm4* and *Npdc1* significantly reduced α-syn PFF-induced neurodegeneration in the SNc (Figure 6a,b, Extended Data Figure 9e) compared to wt PFF-injected mice. As expected, the decrease of TH-positive nigral cells contralateral to the α-syn PFF injection was partial in wt mice, but there was no cell loss in dHET or dKO mice (Extended Data Figure 9e-g). Behavioral analysis of this cohort showed no deficit caused by α-syn PFF in the wt group, so any potential benefit of combined mGluR4 and Npdc1 deletion could not be assessed (Extended Data Figure 9h-k). The protection of nigral DA neurons from α-syn PFF induced neurodegeneration in the transheterozygote state confirms a key role for the interaction of mGluR4 and Npdc1 in synucleinopathy.

**Figure 6.**
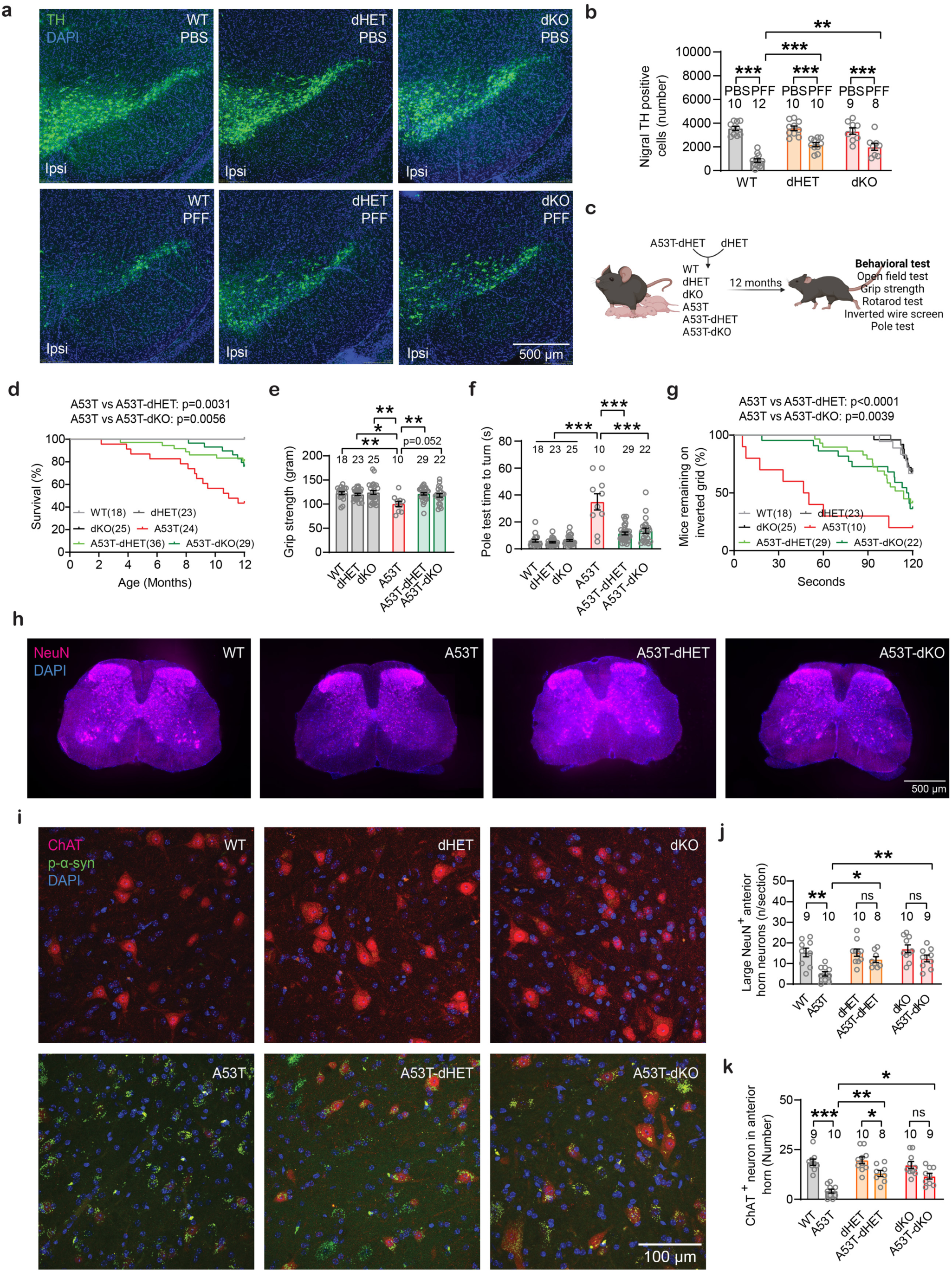
*Npdc1 and Grm4* deletion rescues survival, motor function and neuronal loss induced by α-syn PFF-injection and by A53T expression. **a,** Representative images of TH staining in the ipsilateral side of the SNc of in WT, dHET and dKO mice injected with α-syn PFF. Coronal section with dorsal up and midline to left. Scale bar = 500 µm. **b,** Graph shows mean ±SEM of the number of TH positive neurons in the ipsilateral site of the SNc in WT, dHET and dKO mice injected with α-syn PFF. One-way ANOVA with Tukey’s test. ***p<0.0001 for WT-PBS vs WT-PFF, dHET-PBS vs dHET-PFF, dKO-PBS vs dKO-PFF, and for WT-PFF vs dHET-PPF; **p=0.0030 for WT-PFF vs dKO-PFF. **c,** Diagram illustrating the experimental design for generating and analyzing A53T-dHET and A53T-dKO mice. **d,** Graph shows mean ±SEM of the survival rate in A53T-dHET and A53T-dKO mice. Gehan-Breslow-Wilcoxon test. **e,** Graph shows mean ±SEM of the forelimbs grip strength test in A53T-dHET and A53T-dKO mice. One-way ANOVA with Tukey’s post hoc test. **p=0.0032 for WT vs A53T; *p=0.038 for dHET vs A53T; **p=0.0015 for dKO vs A53T; **p=0.0094 for A53T vs A53T-dHET; *p=0.023 for A53T vs A53T-dKO. **f**, Graph shows mean ±SEM of the time to turn in the pole test in A53T-dHET and A53T-dKO mice. One-way ANOVA with Tukey’s post hoc test. ***p<0.0001. **g,** Graph shows the percentage of mice remaining in the inverted grid in the wire screen test. Gehan-Breslow-Wilcoxon test. **h,** Representative images of NeuN (red) and DAPI (blue) staining of sections L4-L6 of the spinal cord in A53T-dHET and A53T-dKO mice. Scale bar = 500 µm. **i,** Representative images of ChAT (red), p-α-syn (green) and DAPI (blue) staining in the lumbar anterior horn of the spinal cord in A53T-dHET and A53T-dKO mice. Scale bar = 100 µm. **j,** Graph shows mean ±SEM of the number of large NeuN positive neurons per section in the lumbar anterior horn of the spinal cord in A53T-dHET and A53T-dKO mice. One-way ANOVA with Tukey’s post hoc test. **p=0.0020 for WT vs A53T; *p=0.028 for A53T vs A53T-dKO. **k,** Graph shows mean ±SEM of the number of ChAT cells in the lumbar anterior horn of the spinal cord in A53T-dHET and A53T-dKO mice. One-way ANOVA with Tukey’s post hoc test. ***p=0.0006 for WT vs A53T.

## Interaction of *Grm4* and *Npdc1* is required for α-syn A53T transgenic neurodegeneration

To validate these observations in another model, we crossbred *Grm4*-/- and *Npdc1*-/- mice with A53T transgenic mice and generated double heterozygous (dHET) and double ko mice (dKO) on the A53T background (A53T-dHET and A53T-dKO, respectively) as well as wt and A53T littermates (Figure 6c). Body weights were similar among the different genotypes (Extended Data Figure 10a). The A53T mice displayed reduced survival, and the A53T-dHET mice were protected (Figure 6d). Behavioral and histological analysis were performed for surviving mice at 12 months of age. The motor deficit of A53T mice was clear when measuring grip strength, the vertical pole test and the wire screen test (Figure 6e-g). The A53T-dHET and A53T-dKO mice were protected from impairment on each of these motor tasks (Figure 6e-g). No alterations in A53T mice were detected in the open field and the rotarod test (Extended Data Figure 10b-d).

After survival and behavioral studies, histological correlates were examined. We focused on the spinal cord and its motoneurons because they are known to be strongly affected in this strain and are likely responsible for the survival and motor function differences. Astrogliosis as detected by anti-GFAP staining was not prominent in A53T mice as compared to wt and was not altered by the dHET or dKO genotype (Extended Data Figure 10e). In contrast, the A53T mice showed a loss of more than 80% of spinal motoneurons in the ventral horn as compared to wt mice, whether detected as large diameter NeuN positive cells or by staining with choline acetyltransferase (ChAT) (Figure 6h-k). The transheterozygotes and double knockouts for mGluR4 and Npdc1 prevented neuronal loss in the A53T background (A53T-dHET and A53T-dKO mice), even though phosphorylated α-syn accumulated. These results confirm that partial or complete depletion of mGluR4 and Npdc1 rescues the reduced survival rates, motor impairment and neuronal loss in the A53T mice.

## Discussion

We completed a comprehensive expression cloning screen of plasma membrane proteins for high affinity interactions with α-syn PFF. mGluR4 and NPDC1 were identified as α-syn amyloid binding proteins expressed by nigral dopamine neurons. Pathophysiological relevance of these sites for α-syn amyloid triggered disease is supported by multiple lines of evidence. α-syn PFF titrated the physiological function of both mGluR4 and NPDC1. Null mutations of either mGluR4 and NPDC1 prevented nigral dopamine neuron loss triggered by α-syn PFF injection and reduced behavioral deficits associated with PFF injection or mutant α-syn overexpression. Furthermore, the two α-syn binding proteins associated with one another and interacted genetically to mediate α-syn-triggered pathology in both transgenic and fibril injection models. Thus, the mGluR4 – NPDC1 protein complex is a potential therapeutic intervention target for PD, LBD and related disorders.

The observation that loss of cell surface proteins capable of interacting with α-syn amyloid protects nigral dopamine neuron from neurodegeneration supports the hypothesis that synucleinopathy involves a crucial extracellular phase and is not a purely cell-autonomous intracellular process, even in a transgenic overexpression system. This is consistent with a prion or prion-like mechanism of disease progression. However, neurodegeneration does not correlate directly with the accumulation of p-α-syn aggregates^5, 7, 36^. The protection of motoneurons from degeneration triggered by mutant α-syn overexpression and of nigral dopamine neurons from degeneration triggered by striatal PFF injection occurred despite unchanged striatal and spinal cord p-α-syn accumulation. Thus, the presence of mGluR4 and Npdc1 are required for sensitive neurons to detect extracellular aggregates and transduce this information into neuronal loss.

Our studies focused on mGluR4 and NPDC1 because they are expressed by nigral dopamine neurons. Other neuronal binding sites confirmed or identified here include LRRC4C and PRNP. We did not assess their role in neurodegeneration. Both are expressed widely by neurons and may constitute additional neuronal α-syn PFF receptors required for neurodegeneration.

Many of the hits in our screen are expressed primarily or exclusively by microglia and astrocytes. These sites include FCGR1A, FCRL4, SCARF1, LAG3, TRIL, CXCL16, CD72, CD93 and Cdh1. The role of these proteins and glial cells in propagating neuronal inclusions and associated neurodegeneration was not studied here. They may interact with extracellular α-syn amyloids for transfer to neurons, for modification of α-syn protein, for degradation of α-syn, or to stimulate cytokine release. Further study will be required to assess the role of this collection of non-neuronal α-syn PFF binding sites. Given the role of LRRK2 genetic variants in PD risk and the expression of LRRK2 by immune cells^37^, the interaction of LRRK2 signaling with these receptors will be of great interest.

The mGluR4 protein has previously been implicated in modifying basal ganglia function after experimental alteration of nigral DA function. In rodent and non-human primate, acute administration of mGluR4 agonists led to improved motor function acutely and dose dependently^38–41^. This action is distinct from the disease-modifying neuroprotective effect of *Grm4* deletion described here. However, it may be related to the ability of α-syn PFF to modulate mGluR4 signaling and the capacity of mGluR4 agonists to displace α-syn PFF from mGluR4. A mGluR4 positive allosteric modulator has been tested in a phase 2 clinical trial examining short term symptomatic benefit for motor function in PD^26^. While symptomatic benefit did not reach statistical significance the compounds were safe in PD subjects. Any potential disease-modifying effect of mGluR4 pharmacology has not yet been examined.

The presence of low concentrations of α-syn PFF positively modulates L-AP4 activation, while higher concentrations are strongly inhibitory. One interpretation of the biphasic action relates to the dimeric nature of mGluR4; PFF binding to one subunit at low concentrations may differ from effects of binding both subunits at higher concentration. Another possibility is that low α-syn PFF doses may act as an allosteric positive modulator to enhance mGluR4 function. With saturating α-syn PFF concentrations, receptor desensitization may occur, preventing increased activity after subsequent addition with L-AP4.

An unexpected observation was that the two neuronal α-syn binding proteins mGluR4 and NPDC1 physically associate, and that NPDC1 inhibits mGluR4 agonist action. Moreover, the two binding sites interacted genetically to mediate α-syn PFF effects, such that the transheterozygous state greatly reduced α-syn PFF binding, p-α-syn accumulation and synapse loss *in vitro* as well as survival, motor deficits and neurodegeneration *in vivo*. The NPDC1 deletion mapping and the cellular action in *trans* indicate that extracellular contact between mGluR4 and NPDC1 is responsible for the observations. While glutamate is the endogenous ligand for mGluR4, an extracellular polypeptide partner has precedence. The leucine-rich repeat containing transmembrane protein, ELFN1, interacts with mGluR4 and other Class III mGluRs in *trans* with negative allosteric effects^42, 43^.

NPDC1 itself has not been studied in depth previously. In addition to its ability to regulate mGluR4, we observed that NPDC1 overexpressing cells protruded more F-actin rich filopodia. The short cytoplasmic tail of NPDC1 does not provide a clear hypothesis as to how this cellular effect may occur.

Of particular interest for the development of disease-modifying treatment for α-syn-driven neurodegeneration, mGluR4 is an intriguing target. Here, deletion alone or in combination with NPDC1 rescued nigral neurons from striatal α-syn PFF and motoneurons from α-syn-A53T overexpression. Since mGluR4 positive allosteric modulators are known to have some limited symptomatic benefit and we find that they also suppress α-syn amyloid binding to mGluR4, it will be of great interest to optimize mGluR4 pharmacologic to slow or reverse the course of disease driven by misfolded α-syn.

## Author Contributions

Conceptualization, A.P.C, M.C. and S.M.S.; Methodology, A.P.C., M.C., L.A.G., N.K., E.C.G. and S.M.S.; Investigation, A.P.C, M.C., L.A.G., A.H., S.J.T. and N.K.; Writing – Original Draft, A.P.C, M.C. and S.M.S.; Writing – Review & Editing, all; Funding Acquisition, A.P.C. and S.M.S.; Resources, S.M.S; Supervision, S.M.S.

## Acknowledgments

This work was supported by a grant from the Michael J. Fox Foundation to A.P.C., and by grants R01AG034924, R35NS097283, R01AG070926, R01AG066165 and P30AG066508 from the N.I.H. to S.M.S.

## Conflict Of Interest

None.

## METHODS

### α-syn purification and PFF preparation

Untagged α-syn was purified from recombinant bacteria as previously described ^44^. Full-length human α-syn encoded by the pET21a vector (Addgene, #51486) and full-length mouse α-syn encoded by the pET3a vector (Addgene, #108865) were expressed in BL21(DE3) *E. coli (*Invitrogen*, #* C600003). The α-syn monomers were then purified using HiLoad 25/60 Superdex 200 and HR Source Q 20 ml exchange columns (GE Healthcare Life Sciences). Endotoxins were subsequently removed using an endotoxin removal kit (GenScript, #L00338). To generate α-syn PFF (preformed fibrils), the purified α-syn monomers were diluted in phosphate-buffered saline (PBS) to a concentration of 5 µg/µl and agitated for 7 days at 37°C and 1000 rpm to allow the formation of mature fibrils. The aggregates were collected by centrifugation and resuspended in endotoxin-free PBS. These aggregates were then sonicated for 1 minute and 15 seconds (1 second on, 1 second off, 30% amplitude) using Sonifier 250 sonicator (Branson) to generate PFF. The concentration of α-syn was measured using a BCA assay (Thermo Fisher Scientific).

After purification, α-syn was dialyzed in PBS 1x. α-syn fibrils were prepared by incubating the monomeric protein in PBS 1X at 37°C for 7 days in microfuge tubes. The fibrils were stored in aliquots at -80 °C. To generate PFF, α-syn fibrils were sonicated for 1 min 20 sec with 30% power using the Sonifier 250 sonicator (Branson). Sonication was performed right before each experiment. PFF generated from human α-syn monomers were used for experiments in cell lines, while PFF from murine monomers were applied to primary neuronal cultures and stereotaxic injections in mice.

For cell cultures, recombinant α-syn monomer was labeled with sulfo-NHS-LC-Biotin (21335, Thermo Scientific, USA) after purification. Free, unbound biotin was removed from the sample using Amicon Ultra-4 3 kDa tubes (Millipore, #UFC8003). The molar ratio of biotin to α-syn was 2. After biotinylation, α-syn PFF were prepared as mentioned above.

### Atomic Force Microscopy (AFM)

α-syn fibrils and PFF size was analyzed by Atomic Force Microscopy. Briefly, 10 µl of solution were applied to a 10 mm diameter mica discs (Ted Pella, #50) for 5 min, then washed twice with distilled water and let dry before imaging. Imaging was performed using the Dimension FastScan AFM (Bruker) with FASTSCAN-B tips (Bruker) in the tapping mode.

### Expression cloning screening process

HEK293T cells (ATCC) were plated onto Poly-D-Lysine(PDL)-coated 96-well plates (Corning, #354461) in DMEM media (GIBCO, #11965-092) supplemented with 10% fetal bovine serum (GIBCO, #16000-044) and Pen Strep (GIBCO, #15140-122). HEK293T cells were individually transfected with 4,401 human cDNAs encoding membrane proteins from three different libraries. A library from Dharmacon contained 875 cDNAs in pLOC-GFP or pLX304-V5 vector (Dharmacon, #SO-2608946G). The second library was from Transomic and contained 365 cDNA expressing for membrane proteins in pLOC vector (Transomic, #TCH3000). We obtained additional plasmids from a 13,579 cDNA collection in pLX304-V5 plasmid (Transomic, #TOH7500). In order to select clones from this library, we cherry-picked a list of cDNAs classified as plasma membrane encoding genes based on Gene Ontology (GO) classification. Selected cDNAs were re-arranged in 96-well plates (NUNC, #VWR 62407-174). The final number of clones resulting from the 13,579 cDNA collection was 3,161 cDNAs.

Briefly, 15 µl of glycerol stocks for each cDNA were grown in 1 ml of LB media with 100 µg/ml carbenicillin (ref) in deep 96-well plates (USA Scientific, #1896-2110) for 16-18 h at 37°C and 230 rpm. DNA extraction was performed using Plasmid Plus 96 Miniprep Kit (Qiagen, #16181).

HEK293T cells were transfected using Lipofectamine 3000 (ThermoFisher, #L3000015). Two days after transfection, biotinylated α-syn PFF (1000 nM) were added to cell media (DMEM with 10% FBS) and incubated for 2 h at RT. Cells were then washed twice with DMEM media and then with PBS 1x (3 times). Cells were fixed in PFA 4% for 12 min and then blocked for 1 h with BSA 2%, 0.1% Triton X-100 in PBS 1x. Protein expression was monitored with GFP signal or with an anti-V5 antibody (Cell Signaling, #13202S), incubated overnight in blocking solution. PFF binding was detected using Alexa-555 streptavidin (ThermoFisher, #S2138), incubated with a goat anti-rabbit secondary antibody to detect V5 (ThermoFisher, #A-11008) for 1 h. Images were acquired using the automated ImageXpress Micro XLS (Molecular Devices) with a 20x air objective.

Hits obtained were further validated by comparing its binding to biotinylated α-syn PFF with biotinylated monomer and responding to increasing concentrations of α-syn PFF (data not shown).

### NPDC1 deletion mutants

Deletions in human NPDC1 gene in pLX304-V5 plasmid were generated using the Q5 Site-Directed Mutagenesis Kit (NEB, #E0554S) according to the manufacturer’s instructions. The resulting plasmids were transformed into DH5alpha competent cells (Thermofisher Scientific). Integrity of the constructs was verified by sequencing.

### In vitro co-immunoprecipitation

HEK293T cells were transfected with *GRM4*-V5 or *NPDC1*-V5 using Lipofectamine 3000. Two days after transfection, α-syn monomer or PFF at 1000 nM were added to the cells and incubated for 2h at 23°C. Cells were washed twice with media and with PBS 1X and then collected in lysis buffer: 50 mM Tris pH 8.0, 150 mM NaCl, 1% Triton X-100, protease inhibitors cOmplete-mini (Roche, #11836170001). Lysates were centrifuged at 18000 g for 20 min and the supernatant was collected. Protein quantification was determined using Pierce BCA Protein Assay Kit (ThermoFisher, #23225). Then, 500 µg of protein were incubated with 50 µl of protein G magnetic Dynabeads (ThermoFisher, #10003D) conjugated with 5 µg V5 tag monoclonal antibody (ThermoFisher, #R960-25) or mouse IgG2a Isotype Control (CST, #61656). Incubation was performed for 2h at 23°C. Beads were washed 4 times with lysis buffer and eluted with Laemmli buffer at 95°C for 5 min.

### Immunoblot

Protein samples were electrophoresed using precast 4–20% Tris-glycine gels (Bio-Rad, #4561096) and transferred with an iBlot 2 Transfer Device onto nitrocellulose membranes (Invitrogen, #IB23001). The membranes were incubated in blocking buffer (Rockland, #MB-070) for 1 h at room temperature and then incubated overnight at 4°C in blocking buffer with primary antibody against V5-Tag D3H8Q (CST, 13202). The next day, membranes were washed three times with TBST for 3 min and incubated in secondary antibodies (Li-Cor, IR Dye 680 or 800, 1:10,000) for 1 h at room temperature. After washing three times with TBST for 3 min, proteins were visualized with an Odyssey Infrared imaging system (Li-Cor). The immunoreactive bands were quantified using ImageJ software.

### Immunofluorescence of cultures

HEK293T cells and cultured neurons were washed twice with cell media and then 3 times with PBS1x. Cells were fixed in PFA 4% for 12 min and then blocked for 1 h with BSA 2%, 0.1% Triton X-100 in PBS 1x. Cells were then incubated overnight with primary antibodies in blocking buffer at 4°C: V5 antibody (Cell Signaling, #13202S, 1:1000) or MAP2 (Millipore, # AB5622, 1:1000). The cells were then washed 3 times with DPBS and incubated for 2 h at room temperature in Alexa Fluor 488 donkey anti-rabbit IgG (H + L) (Invitrogen #A21206, 1:1000) in blocking buffer. Biotinylated α-syn PFF binding was detected using streptavidin-555 (ThermoFisher, #S2138, 1:1000). Images were acquired using the automated ImageXpress Micro XLS (Molecular Devices) with a 20x air objective for HEK293T cells. Stained neurons were imaged using LSM800 confocal microscopy (Zeiss) with 40x magnification objective lens.

For automated microscopy of HEK293T cells and primary neurons using ImageXpress® Micro Confocal system, the cells were fixed with 4% paraformaldehyde for 15 minutes at 4°C, followed by blocking and permeabilization using a solution of 10% horse serum and 0.1% Triton X-100 for 1 hour at room temperature. Subsequently, the cells were incubated overnight at 4°C in a primary antibody solution consisting of 1% normal horse serum, 0.1% Triton X-100 in DPBS. The following primary antibodies were used: anti-V5 (CST 80076S, 1:500), anti-DYKDDDDK Tag (CST 14793S, 1:500), anti-MAP2 (Abcam AB5392, 1:10000), anti-p-α-syn (Invitrogen PA14686, 1:2000), anti-Synapsin1/2 (Synaptic System 106004, 1:500), and anti-Homer 1b/c (Synaptic System 160111, 1:500). After primary antibody incubation, cells were washed three times with DPBS, then incubated with fluorescent secondary antibodies (Invitrogen Alexa Fluor 1:500) in 1% normal donkey serum and 0.2% Triton X-100 in DPBS for 1-2 hours at 23°C.

For single V5 staining, Alexa Fluor 488 (Invitrogen # A-21202) was used. For V5/Flag co-staining, Alexa Fluor 488 (Invitrogen # A-21202) and Alexa Fluor 647 (Invitrogen # A-31573) were applied. For α-syn-PFF/MAP2 co-staining, Alexa Fluor 555 streptavidin (Invitrogen # S21381) and Alexa Fluor 647 (Invitrogen # A-78952) were used. In the case of p-α-syn /MAP2 co-staining, Alexa Fluor 488 (Invitrogen # A-11008) and Alexa Fluor 647 (Invitrogen # A-78952) were used. For triple staining of Synapsin1/2, Homer1, and MAP2, the secondary antibodies applied were Alexa Fluor 488 (Invitrogen # A-11001), Alexa Fluor 555 (Invitrogen # A-21435), and Alexa Fluor 647 (Invitrogen # A-21450), respectively.

After secondary antibody incubation, nuclei were stained with DAPI (Invitrogen D1306, 1 µg/mL in DPBS) for 15 minutes at room temperature, followed by three washes with DPBS. Images of HEK293T cells and primary neurons were captured using the ImageXpress® Micro Confocal system (Molecular Devices) with 20X or 40X objective lenses.

### mGluR4 signaling assays

α-syn PFF binding assays in the presence of mGluR4 modulators were conducted in HEK293T cells. Cells were plated onto PDL-coated 96-well plates (Corning, #354461) in DMEM media (GIBCO, #11965-092) supplemented with 10% fetal bovine serum (GIBCO, #16000-044) and Pen Strep (GIBCO, #15140-122). Cells were transfected with GRM4-V5 in a pLX304 plasmid using Lipofectamine 3000 (ThermoFisher, #L3000015). Two days after transfection, media was replaced by DMEM without FBS 30 min before the experiment. Then, mGluR4 modulators were added for 30 min at RT: glutamate (Sigma, #G1626), L-AP4 (Abcam, #ab144481), and MSOP (Tocris, #0103). Cells were then treated with biotinylated α-syn PFF for 2h at RT. Immunostaining was performed as previously described. Images were acquired using the automated ImageXpress Micro XLS (Molecular Devices) with a 20x air objective

HEK293T cells (passages 5 to 20) were cultured in DMEM GlutaMAX (GIBCO, #10566-016) supplemented with 10% FBS (Seradigm, #1500-500) and 2 mM sodium pyruvate (ThermoFisher Scientific, #11360070) at 37°C with 5% CO2. mGluR4 Gαqi5 stable HEK293T cells (passages 3 to 20, Multispan, #CG1191) were cultured in DMEM GlutaMAX with 10% dialyzed FBS (ThermoFisher Scientific, #26400044), 2 mM sodium pyruvate, 1 μg/ml puromycin (Invivogen, #ant-pr-5), and 250 μg/ml hygromycin (Invivogen, #ant-hm-5). Cells were transiently transfected with either a synthetic vector encoding pcDNA3.1 or human NPDC1 cDNA in the pLX304 vector using Lipofectamine 3000 (ThermoFisher Scientific, #L3000-008) according to the manufacturer’s protocol.

For experiments involving only mGluR4 Gαqi5 stable HEK293T cells or transfected mGluR4 Gαqi5 stable HEK293T cells, cells were plated onto PDL-coated 96-well Black/Clear Flat Bottom TC-treated Assay Plates (Corning, #354640) at a density of 1x10^5^ cells/ml. HEK293T cells were transfected in 6-well plates for 24 hours, followed by a media change to DMEM without L-glutamine (HyClone, #SH30285.01) supplemented with 10% dialyzed FBS, and incubated at 37°C with 5% CO_2_ for an additional 24 hours. The medium was then collected as conditioned medium. Transfected cells were resuspended and mixed with mGluR4 Gαqi5 stable HEK293T cells at a ratio of either 1:1 or 2:1 (maintaining 1x10^4^ cells per well for Grm4-HEK293T cells) using DMEM without L-glutamine (10% dialyzed FBS), and plated onto PDL-coated 96-well plates. They were incubated at 37°C for another 24 hours.

For experiments using only mGluR4 Gαqi5 stable HEK293T cells, the medium was changed to DMEM without L-glutamine (10% dialyzed FBS) one day before the FLIPR experiment. Prior to the conditioned medium assay, the medium was replaced with conditioned medium and incubated at 37°C with 5% CO_2_ for 2 hours. On the day of the FLIPR experiment, FLIPR Calcium 6 dye was diluted with Hanks’ Balanced Salt Solution (HBSS) supplemented with 5 mM Probenecid (Invitrogen, #P36400). Then, 100 µL of the diluted dye was added to each well, and the plate was incubated at 37°C with 5% CO2 for 2 hours. Fluorescence measurements were taken using the FlexStation 2 (Molecular Devices) over 120 seconds at 2-second intervals. 20 µL of L-AP4 or DPBS was injected into the wells at 20 seconds, followed by a similar injection program for Ionomycin (Sigma, #I0634-1MG). The relative fluorescence units (RFU) were calculated by subtracting the basal signal (mean signal from 0 to 20 seconds) from the raw signal (mean signal from 20 to 180 seconds).

### Npdc1 modulation of F-Actin

HEK293T cells were plated onto 8-well Lab-Tek chamber slides (Nunc, #154941PK) in DMEM media (GIBCO, #11965-092) supplemented with 10% fetal bovine serum (GIBCO, #16000-044) and Pen Strep (GIBCO, #15140-122). Cells were transfected with *NPDC1*-V5 in a pLX304 plasmid using Lipofectamine 3000 (ThermoFisher, #L3000015). Two days after transfection, cells were treated with biotinylated α-syn PFF for 3h at 37°C. Cells were then washed twice with media and 3 times with PBS 1x. Immunofluorescence was performed as previously indicated. Alexa-555 Phalloidin was used to detect actin filaments (ThermoFisher, #A34055) and α-syn PFF were detected with Alexa-488 streptavidin (ThermoFisher, #S32354) in blocking buffer.

### Neuronal culture and PFF exposure

Cortical cells were isolated from the cortices of E16-17 mouse embryos as reported previously^45^. The embryos were decapitated, and the anterior cortex was dissected out under a stereotactic microscope using BrainBits Hibernate E minus calcium medium, kept on ice. The tissue was then digested with a freshly prepared enzyme solution (Mg/Ca-free HBSS containing 20 U/ml Papain (Worthington LK003178), 3 mM EDTA (AmericanBio), 2 mM CaCl₂ (VWR E506), and 1 mg/ml DNAse (Sigma DN25)) at 37°C for 30 minutes. After digestion, the cell suspension was filtered through a 40 µm cell strainer, and the cells were counted and diluted to a concentration of 1x10⁵ cells/ml. A 100 µl volume of this suspension was plated onto a PDL-coated 96-well Black/Clear Flat Bottom Plate. The plates were incubated at 37°C with 5% CO₂, and the medium was half-changed every 7 days.

For the binding assay, PFF was added at DIV 17-18 and incubated at 4°C or 37°C for 2 hours. For the α-syn phosphorylation and synaptic loss assays, PFF were added at DIV 10 and incubated at 37°C for 7 days.

### Mice

All protocols were approved by Yale Institutional Animal Care and Use Committee (IACUC) (2024-07281). Npdc1 knock-out mouse embryos were obtained from EMMA (B6;129X1-Npdc1^tm1Ce^/Orl, #EM:00211) and rederived at Yale Genome Editing Center. Grm4 knock-out mice with C57BL/6 J background (B6.129-Grm4^tm1Hpn^/J, #003619) and human α-syn A53T transgenic mice with C57BL/6 J background (B6.Cg-2310039L15Rik^Tg(Prnp-SNCA*A53T)23Mkle^/J, #006823) were obtained from the Jackson Laboratory. The mice were grouped and housed 2– 5 per cage, adhering to a 12-hour light-dark cycle, with constant access to food and water. The room conditions were maintained at a temperature of 21–23°C and a humidity of 50±20%.

### α-syn PFF stereotaxic injections

Immediately before the intrastriatal injections, fibrils were diluted in sterile PBS and sonicated as indicated above. PBS 1X or recombinant α-syn PFF (5 µg in 2 µl) was stereotactically delivered into the right dorsal striatum using the following coordinates: +0.2 mm Medial-lateral (ML); +2.0 mm antero-posterior (AP) and +2.8 mm dorso-ventral (DV) from bregma. Injections were performed using a 10 µL syringe (Hamilton, #7635-01) and 33-gauge needle, 45° tip (Hamilton, #7803-05) at a rate of 0.1 µl per minute. After delivery, the needle was left in place for 5 minutes and then withdrawn over 2 minutes. After surgery, mice received 0.05 mg/kg buprenorphine for 3 days (twice daily, 12 h apart) as analgesic and body weight was monitored once a month.

Animal behavior was performed after 6 months and mice were then euthanized. For histological studies, mice were perfused transcardially with PBS and 4% PFA and brains were removed, followed by overnight fixation in 4% PFA and washed and kept in PBS 1X after.

### Immunohistology of brain sections

All histological tests were completed and analyzed by experimenter blind to genotype and group.

Mice were euthanized via CO2 inhalation, followed by perfusion with ice-cold DPBS and 4% paraformaldehyde in sequence. The brains and spinal cords were quickly dissected and sectioned into 40 μm slices using a Leica WT1000S vibratome. The sections were blocked and permeabilized in 10% normal horse serum with 0.1% Triton X-100 for 1 hour at room temperature. Following this, the sections were incubated overnight at 4°C with primary antibodies diluted in 1% normal horse serum, 0.1% Triton X-100 in DPBS. The primary antibodies used were: anti-TH (EMD Millipore AB152, 1:500), anti-p-α-syn (BioLegend 825701, 1:500), anti-NeuN (EMD Millipore ABN91, 1:500), anti-ChAT (EMD Millipore AB144P, 1:100), and anti-GFAP (DAKO Z0334, 1:500).

After incubation, the sections were washed three times with DPBS and incubated for 1-2 hours at room temperature with fluorescent secondary antibodies (Invitrogen Alexa Fluor, 1:500) in 1% normal donkey serum and 0.2% Triton X-100 in DPBS. For TH single staining, Alexa Fluor 488 (Invitrogen # A-11008) was used. For p-α-syn/TH co-staining, Alexa Fluor 488 (Invitrogen # A-11008) and Alexa Fluor 647 (Invitrogen # A-31571) were used. For NeuN/GFAP co-staining, Alexa Fluor 488 (Invitrogen # A-11008) and Alexa Fluor 647 (Invitrogen # A-78952) were employed. For ChAT/GFAP/p-α-syn triple staining, Alexa Fluor 488 (Invitrogen # A-11008), Alexa Fluor 555 (Invitrogen # A-21432), and Alexa Fluor 647 (Invitrogen # A-31571) were used. Following antibody incubation, nuclei were stained with DAPI (Invitrogen D1306, 1 µg/mL in DPBS) for 15 minutes at room temperature, followed by three DPBS washes. The sections were mounted on glass slides and coverslipped using Vectashield antifade mounting medium (Vector).

Images of TH single staining in nigra sections and NeuN/GFAP co-staining in spinal cord sections were captured using an Olympus VS200 Slide Scanner with a 10X objective lens. Confocal images of p-α-syn/TH co-staining and ChAT/GFAP/p-α-syn triple staining were obtained with a Leica DMi8 confocal microscope using a 40X objective lens.

### Image quantification of immunohistochemistry

All images were obtained, processed, and analyzed by a researcher who was unaware of the animal genotypes and group information.

All quantitative image analyses conducted with the Leica DMi8 confocal microscope were processed using ImageJ (National Institutes of Health). For the p-α-Syn/TH co-staining of brain slices, we applied uniform thresholding and binarization across all images, followed by measuring the areas of p-α-Syn and TH. In the ChAT/GFAP/p-α-Syn triple staining of spinal cord slices, we also applied uniform thresholding and binarization, manually counted the ChAT-positive cells, and measured the GFAP intensity in the anterior horn.

Similarly, all quantitative image analyses from the Olympus VS200 Slide Scanner microscope were processed using ImageJ. To evaluate neuronal loss in the substantia nigra stereologically, we selected every sixth serial coronal slice of 40 µm thickness through the entire substantia nigra, applied uniform thresholding and binarization, and counted all TH-positive cells. From this standardized sampling method, we calculated the DA neuron count per SNc. For NeuN staining, we manually counted the larger NeuN-positive cells in the anterior horn.

Quantitative analyses of images acquired using the ImageXpress® Micro Confocal system were processed with MetaXpress software. After uniformly thresholding and binarizing the images, we measured MAP2 intensity and then expanded the MAP2 signal by one pixel using the “Grow Objects” function. The MAP2 signal was set as a mask to measure streptavidin (PFF) intensity on MAP2 for PFF/MAP2 co-staining, and p-α-Syn intensity on MAP2 for p-α-Syn/MAP2 co-staining. Additionally, we quantified the number of “active synapses” on MAP2 by using the “Found Round Objects” tool to detect Synapsin1/2 and Homer1b/c double-positive puncta on MAP2, with diameters ranging from 0.5–2 µm and intensities of 1500 above local background.

### Behavioral tests

All behavioral tests were performed and analyzed by experimenter blind to genotype and group.

#### Pole test

It was performed with a wooden pole of 57 cm in height and 2 cm in width. Animals were acclimatized for two consecutive days prior to the test and each time five trials were given to each animal. During the test, animal was placed near the top of the pole facing upwards. The time taken by each animal to turn downwards was monitored as the pole turn time. Moreover, the time taken by each animal to reach the base of the pole was recorded as the pole climb downtime. The surface of the pole was made rough by covering it with bandage gauze. The maximum time for recording was set as 60 s. If any animal was found to stall for more than 60 s, the test was further performed for that particular animal.

#### Inverted wire screen test

The inverted wire screen test of motor function was performed by placing the mouse upright on a platform covered with a wire screen and then inverting the platform. The latency of the mouse to fall off of the wire screen was measured, and the better time was taken from two trials. Trials were stopped if a mouse reached a criterion of 120 s.

#### Open field

Open field test was conducted to look at the locomotor activities of the animals. Locomotor activity was monitored with a camera linked to Noldus system and EthoVisionXT software (Netherlands). The instrument records the overall movement abilities of the animals including total distance moved, velocity, total moving time, resting time, center time, and frequencies of movement and rest. Before the test, mice were placed inside the open field arena for 10 min daily for two consecutive days to train them and record their baseline values. Two days after the training, each mouse was taken from the cage and gently placed in the middle of the open field arena. After releasing the animal, data acquisition was started by the software for the next 5 min and the parameters related to the locomotor activities were collected by the software.

#### Rotarod

Prior to the rotarod test, mice were placed on the rotarod instrument for 5–10 min daily for consecutive two days to train them. After 2 days of training, mice were placed on the rotating rod, which rotates with a gradual increasing speed of 4–40 rpm. The experiment was ended if the animal slips from the rotating rod to the base of the instrument or just grips the rod to turn reverse without rotating against the direction of rotating rod.

#### Grip strength test

Grip strength was measured using a grip force meter from Columbus Instruments (Columbus, OH, USA). During each assessment, mice were placed on the apparatus and gently pulled parallel to the ground until they could no longer maintain their grip. Each mouse underwent five trials at every time point, with a 10-second rest period between each trial. The highest three scores from the five trials were averaged to calculate the final grip strength score for each mouse.

### Statistical analysis

Data are shown as mean ± SEM. One-way ANOVA with Tukey’s correction (for multiple comparisons), Gehan-Breslow-Wilcoxon test, two-way ANOVA with Sidak’s post hoc test, and Kruskal-Wallis test with Dunn’s post hoc test were analyzed using Prism (version 9.0.1) software. Gaussian distributions were assumed based on the previous studies. Separate dots on each figure correspond to *N* values reflecting different mice. Data are reported as statistically significant for p < 0.05 (*p < 0.05; **p < 0.005; *** < 0.001).

## Extended Data

**Extended Data Figure 1.**
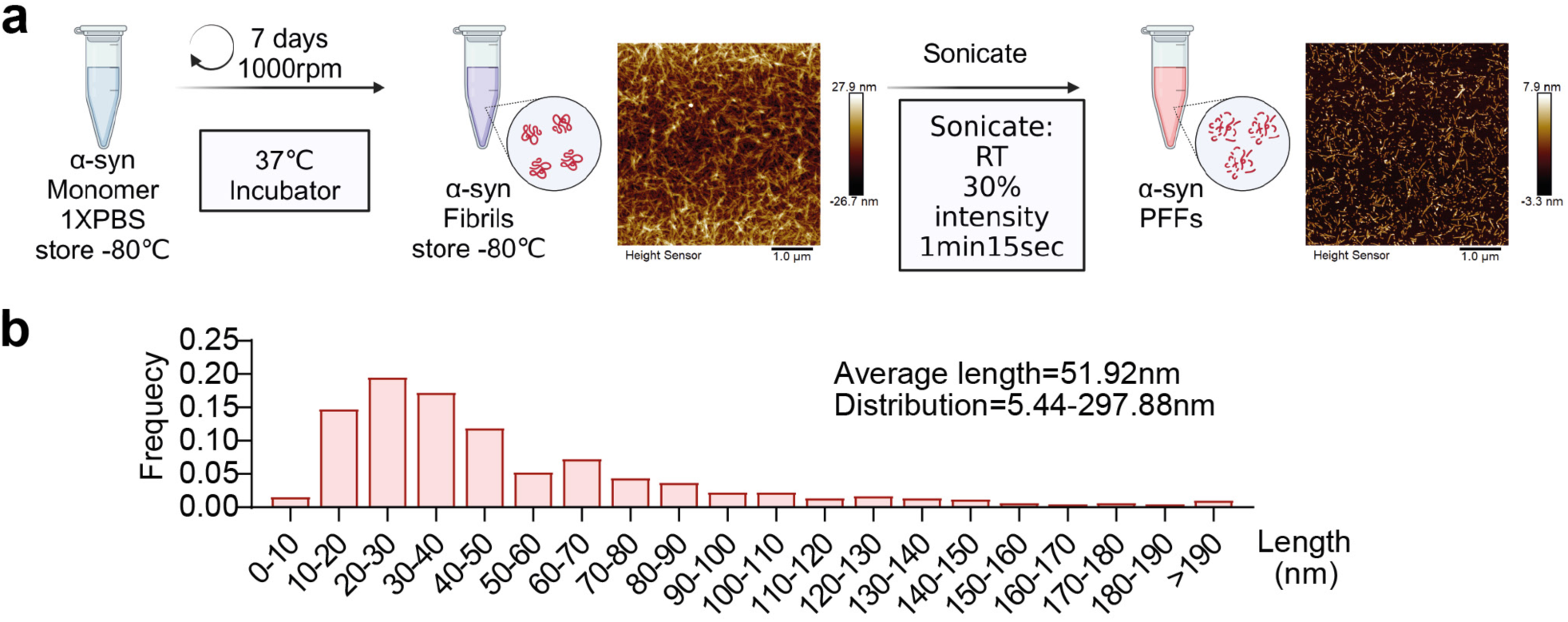
Preparation of α-syn Pre-Formed Fibrils (PFF). **a,** Diagram illustrating the methodology for α-syn PFF preparation. All details are summarized in the Methods section. **b,** Graph shows the frequency of α-syn PFF particles of different lengths as determined by AFM.

**Extended Data Figure 2.**
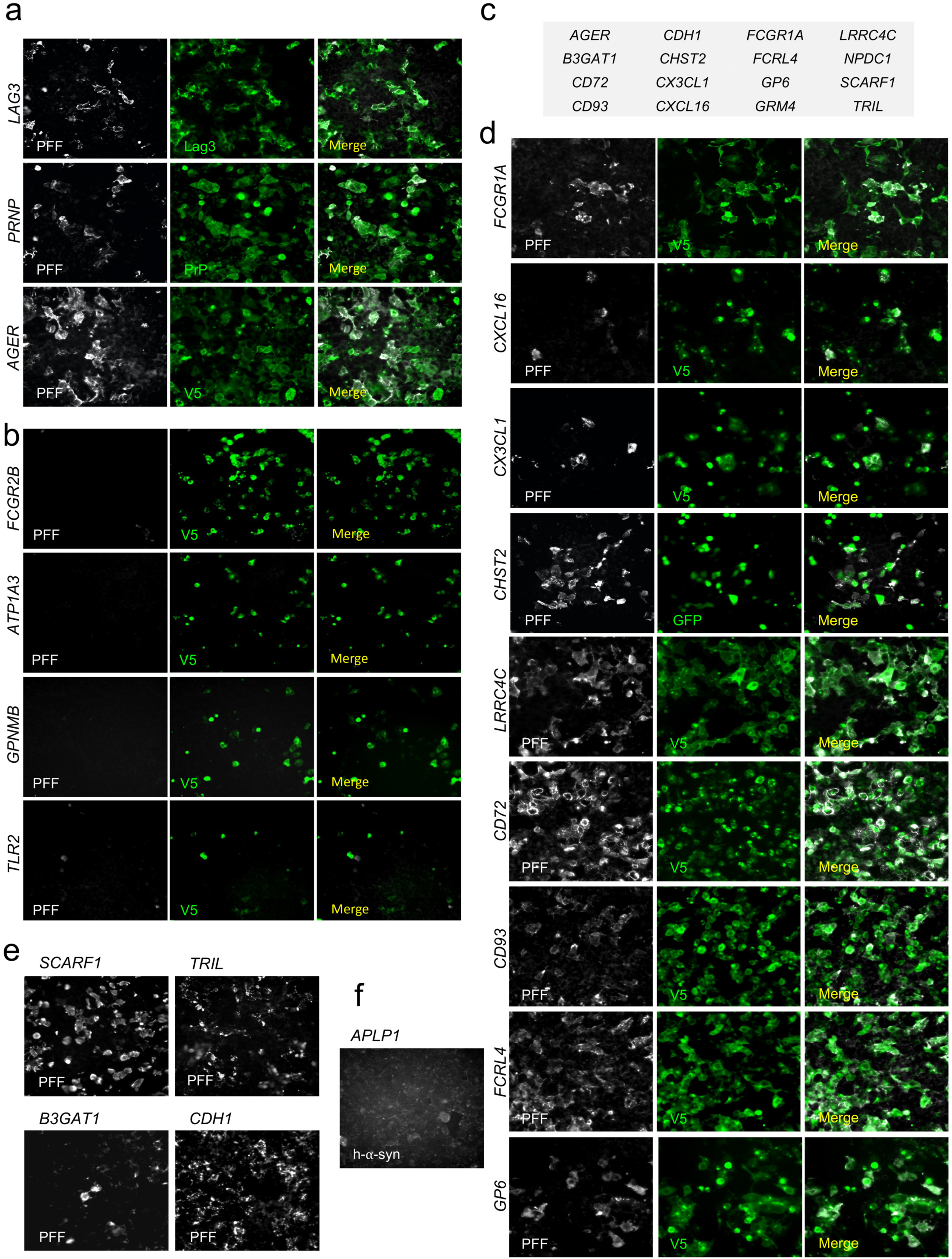
Expression screening for α-syn PFF cell surface binding proteins. Representative images of HEK293T cells transfected with different cDNAs and treated with biotinylated α-syn PFF (1000 nM) for 2 h and stained for PFF binding (streptavidin, gray) and protein expression (green). **a,** HEK293T cells overexpressing *LAG3, PRNP* and *AGER*, which were previously described as α-syn PFF interactors and which detected PFF binding here. **b,** HEK293T cells transfected with *FCGR2B, ATP1A3* (N^+^/K^+^-ATPase), *GPNMB* and *TLR2*, which were described in the literature as α-syn PFF receptors but were negative for α-syn PFF binding under these conditions. **c,** List of confirmed cDNA expression vectors supporting high affinity α-syn PFF cell surface binding when transfected into HEK293T cells. **d, e,** HEK293T cells overexpress V5 tagged (**d**) or untagged (**e**) novel proteins found in our screening as α-syn PFF interactors. **f,** *APLP1*, previously described as PFF receptor, expressed in HEK293T cells treated with non-biotinylated α-syn PFF. In this case, PFF binding was examined with a human specific α-syn antibody.

**Extended Data Figure 3.**
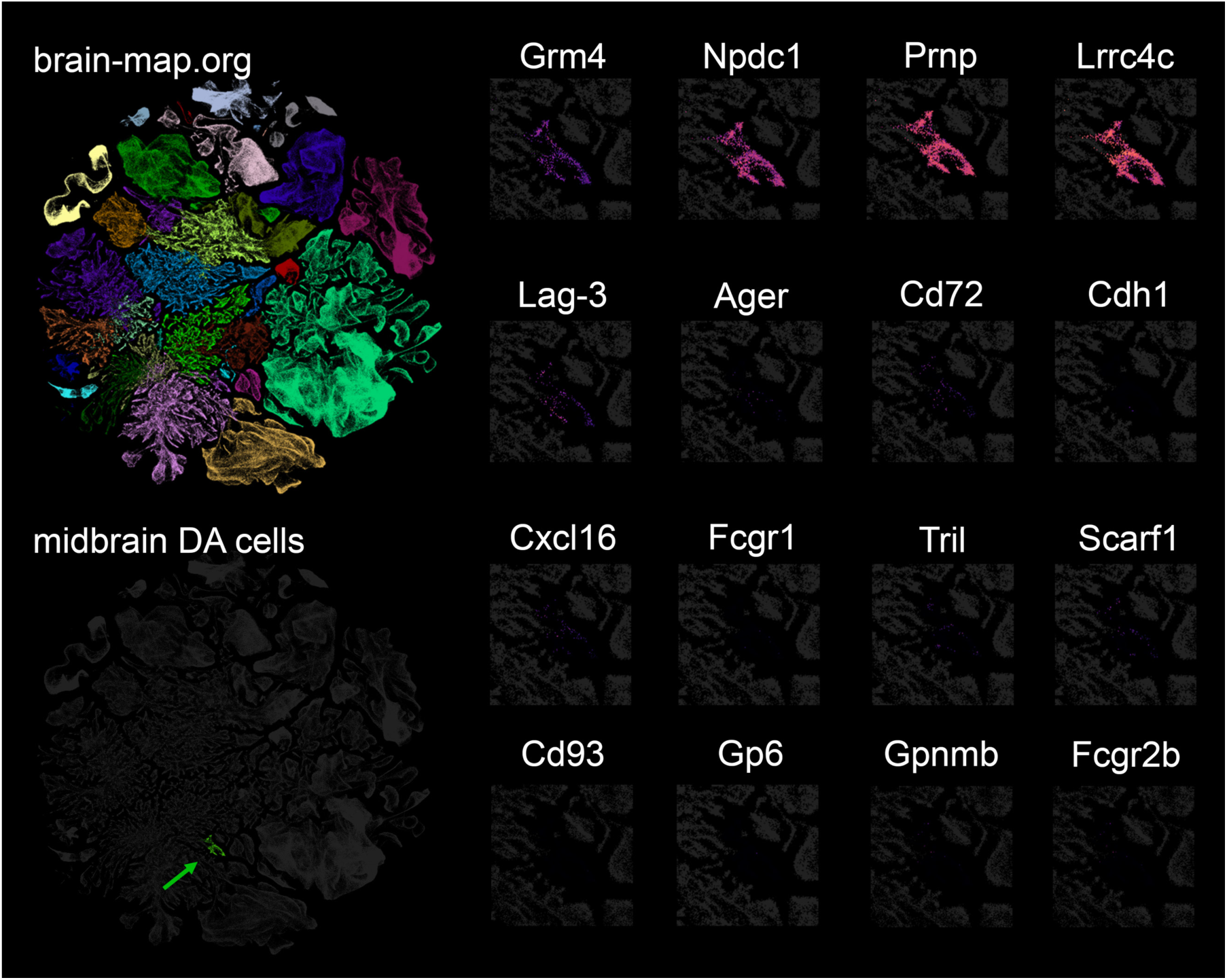
Expression of α-syn PFF binding proteins in mouse midbrain dopamine cells. Expression data were accessed through the brain-map.org website (all clusters of nuclei at top left), and filtered to show only expression in midbrain dopamine cells (green arrow in bottom left). The individual panels on the right show single cell expression of the indicated genes in a magnified view of nuclei in the expression space containing midbrain DA cells. The purple-red color intensity reflects the mRNA counts per million reads in the midbrain DA cluster for each of the listed genes. Note positive signal for *Grm4*, *Npdc1*, *Prnp* and *Lrrc4c*.

**Extended Data Figure 4.**
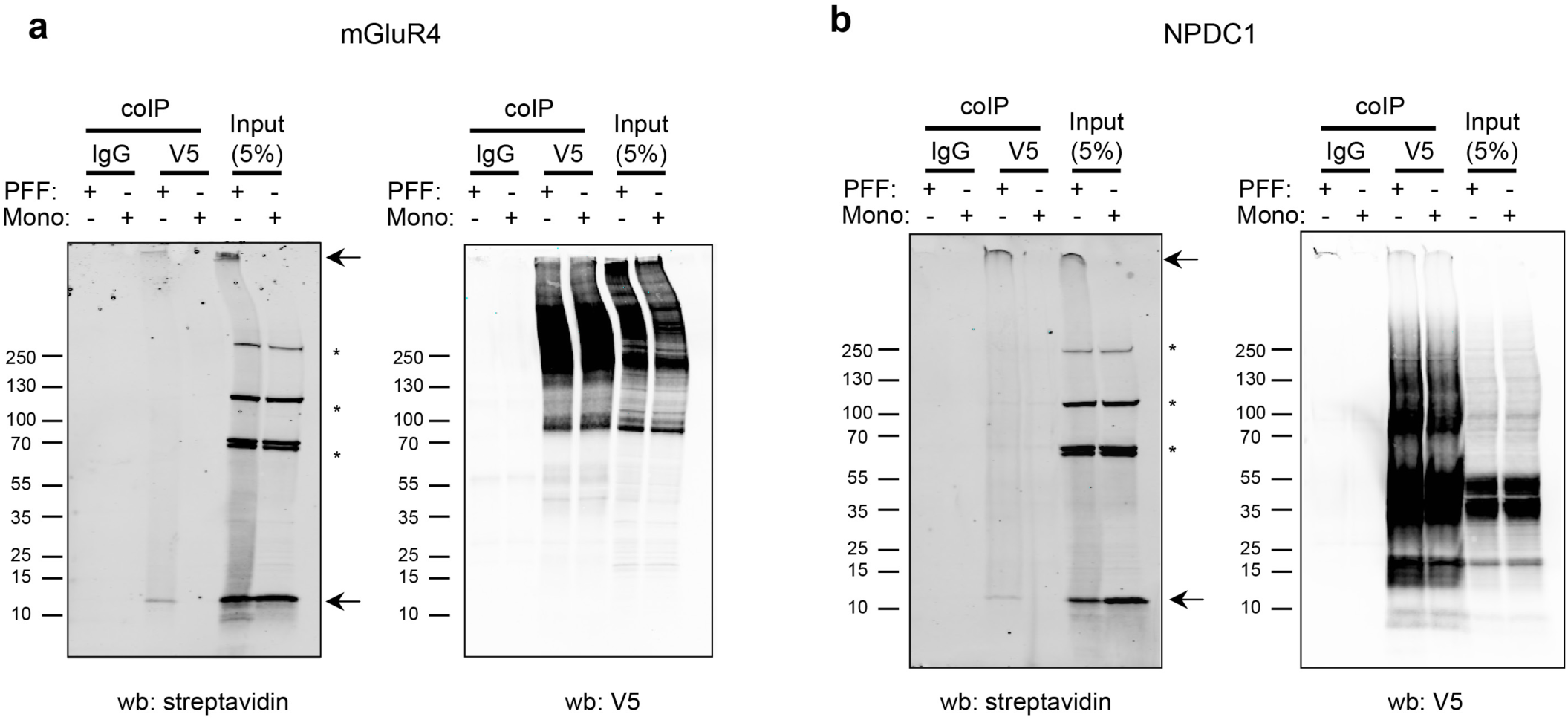
α-syn PFF co-immunoprecipitation with GMR4 and NPDC1. Representative immunoblots after co-IP assay in HEK293T cells with anti-streptavidin (PFF) or V5 (**a**: *GRM4*; **b**: *NPDC1*). Arrows indicate biotinylated α-syn PFF bands in the streptavidin blot. Asterisks indicate non-specific immunoglobulin bands in the streptavidin blot.

**Extended Data Figure 5.**
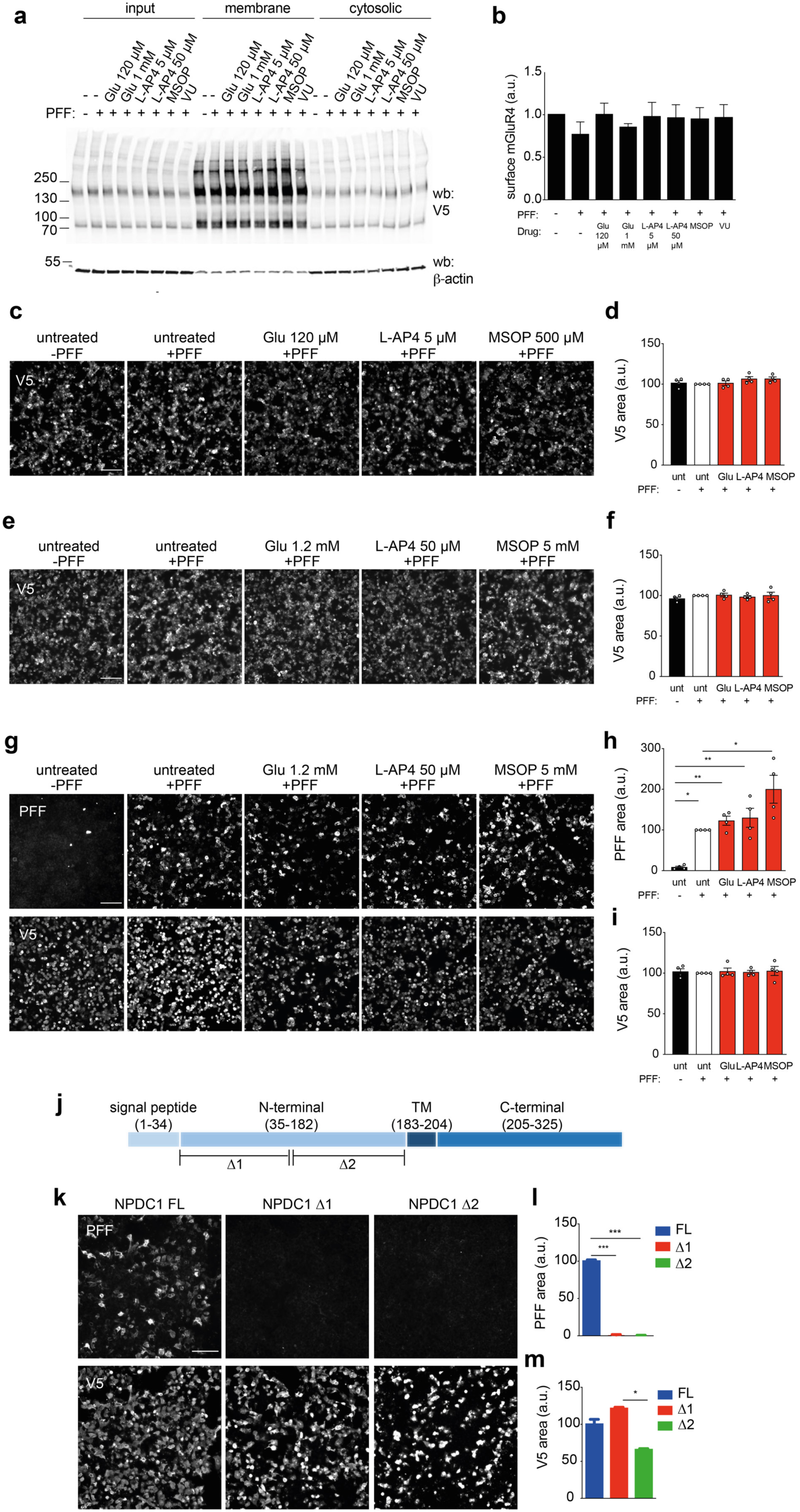
mGluR4 ligands do not alter protein expression and levels at the plasma membrane while NPDC1 deletions block α-syn PFF binding. **a,** Representative immunoblot with anti-V5 in HEK293T cells transfected with *GRM4* and treated with different mGluR4 ligands (glutamate, L-AP4, MSOP and VU). Surface biotinylation was performed in living cells to collect the plasma membrane fraction. **b,** Graph shows surface levels of mGluR4 in HEK293T cells after treatment with the different compounds. One-way ANOVA with Tukey’s post hoc test. **c,** Representative images of HEK293T cells transfected with *GRM4* and treated with mGluR4 ligands (Glu, L-AP4, MSOP) before the addition of biotinylated α-syn PFF for 1 h. Cells were stained with V5 to detect mGluR4 expression. **d,** Graph shows mean ± SEM of the area occupied by V5 in HEK293T treated with biotinylated α-syn PFF for 1 h. One-way ANOVA with Tukey’s post hoc test. **e,** Representative images of HEK293T cells transfected with *GRM4* and treated with mGluR4 ligands (Glu, L-AP4, MSOP) before the addition of biotinylated α-syn PFF for 3 h. Cells were stained with V5 to detect GRM4 expression. **f,** Graph shows mean ± SEM of the area occupied by V5 in HEK293T treated with biotinylated α-syn PFF for 3 h. One-way ANOVA with Tukey’s post hoc test. **g,** Representative images of HEK293T cells transfected with *NPDC1* and treated with mGluR4 ligands (Glu, L-AP4, MSOP) before the addition of biotinylated α-syn PFF for 3 h. PFF bound to the cells are detected with streptavidin. Cells were stained with V5 to detect NPDC1 expression. **h,** Graph shows mean ± SEM of the area occupied with α-syn PFF in HEK293T treated with biotinylated α-syn PFF for 3 h. One-way ANOVA with Tukey’s post hoc test. **i,** Graph shows mean ± SEM of the area occupied by V5 in HEK293T treated with biotinylated PFF for 3 h. One-Way ANOVA with Tukey’s post hoc test. **j,** Schematic representation of NPDC1 protein structure indicating the deletion sites in the extracellular portion. **k,** Representative images of HEK293T cells transfected with *NPDC1-FL, NPDC1-Δ1* and *NPDC1-Δ2* and treated with biotinylated α-syn PFF for 2h at RT. α-syn PFF binding was detected with streptavidin. Cells were stained with V5 to detect NPDC1 expression. **l,** Graph shows mean ± SEM of the area occupied with α-syn PFF in HEK293T treated with biotinylated α-syn PFF for 2 h. One-way ANOVA with Tukey’s post hoc test. **m,** Graph shows mean ± SEM of the area occupied with V5 in HEK293T treated with biotinylated α-syn PFF for 2 h. One-way ANOVA with Tukey’s post hoc test.

**Extended Data Figure 6.**
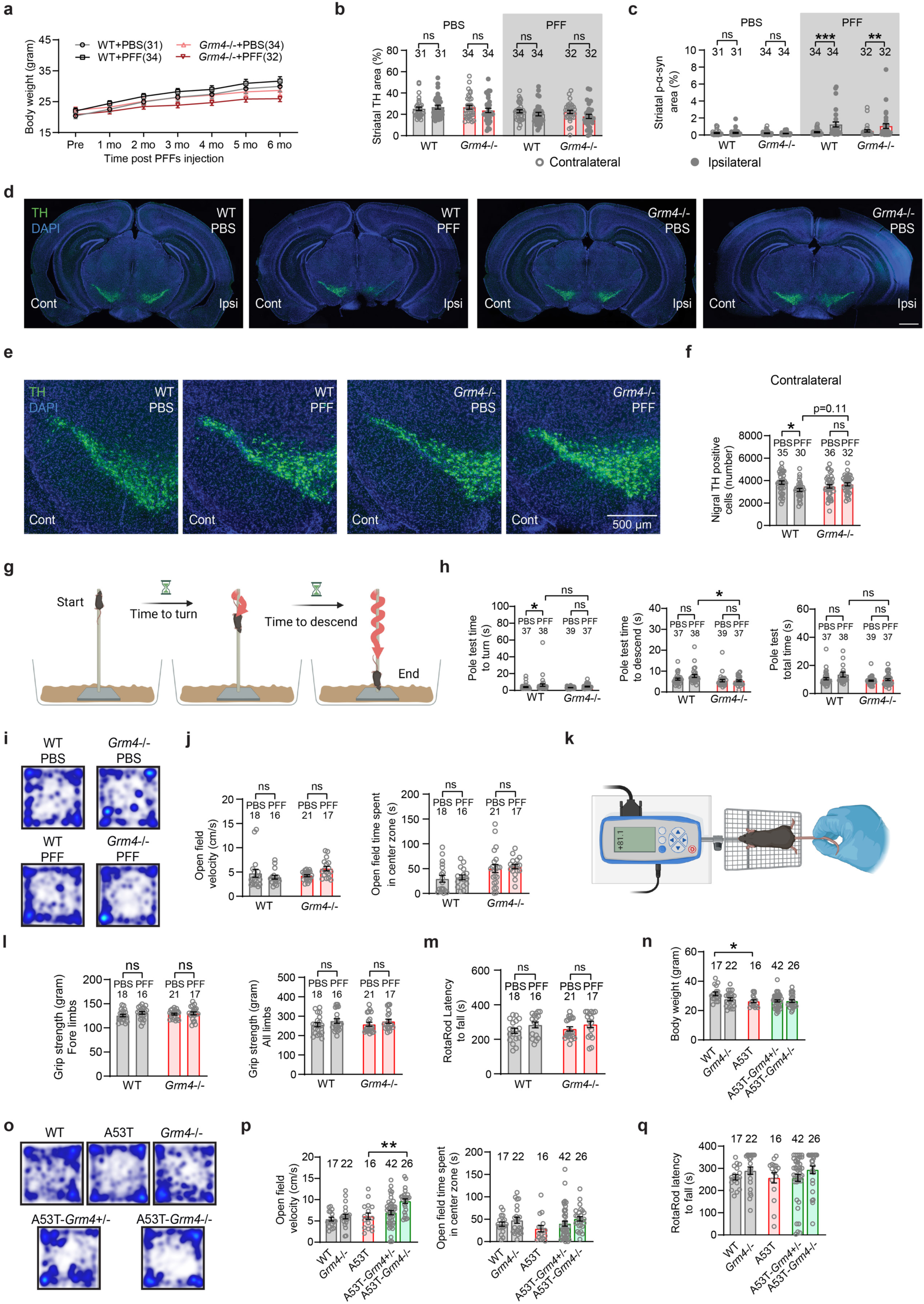
Additional phenotypic measures in *Grm4*-/- mice injected with α-syn PFF and in A53T-*Grm4*-/- transgenic mice. **a,** Graph shows mean ± SEM of the body weight in *Grm4*-/- mice injected with α-syn PFF. **b,** Graph shows mean ± SEM of the TH area in the ipsilateral and contralateral sides of the SNc in *Grm4*-/- mice injected with α-syn PFF. Two-way ANOVA with Sidak’s post hoc test. **c,** Graph shows mean ± SEM of the p-α-syn area in the ipsilateral and contralateral sides of the SNc in *Grm4*-/- mice injected with α-syn PFF. Two-way ANOVA with Sidak’s post hoc test. **p<0.0001 for WT-PFF contralateral vs ipsilateral and **p=0.0028 for Grm4 KO-PFF contralateral vs ipsilateral. **d, e,** Representative images of TH (green) and DAPI (blue) staining in (**d**) the entire brain and (**e**) contralateral side of the SNc in *Grm4*-/- mice injected with α-syn PFF. In **e**, coronal section with dorsal up and midline to right. Scale bars = 500 µm. **f,** Graph shows mean ± SEM of the number of TH positive neurons in *Grm4*-/- mice injected with α-syn PFF. One-way ANOVA with Tukey’s post hoc test. *p=0.018 for WT-PBS vs WT-PFF. **g,** Diagram illustrating the pole test performance. **h,** Graphs show mean ± SEM of the time to turn (left), time to descend (center) and total (right) time in the pole test in *Grm4*-/- mice injected with α-syn PFF. **i,** Representative images of the open field performance in *Grm4*-/- mice injected with α-syn PFF. **j,** Graphs show mean ± SEM of the velocity (left) and the time spent in the center (right) in the open field test. One-way ANOVA with Tukey’s post hoc test?? **k**, Diagram illustrating the grip strength measurement. **l**, Graphs show mean ± SEM of the grip strength of the forelimbs (left) and all limbs (right) in *Grm4*-/- mice injected with α-syn PFF. One-way ANOVA with Tukey’s post hoc test. **m**, Graph shows mean ± SEM of the latency to fall in the rotarod test in *Grm4*-/- mice injected with α-syn PFF. One-way ANOVA with Tukey’s post hoc test. **n**, Graph shows mean ± SEM of the body weight in *Grm4*-/- mice injected with α-syn PFF. One-way ANOVA with Tukey’s post hoc test. **p=0.025 for WT vs A53T mice. **o,** Representative images of the open field performance in A53T-*Grm4*-/- mice. **p,** Graphs show mean ± SEM of the velocity (left) and the time spent in the center (right) in the open field test. One-way ANOVA with Tukey’s post hoc test. For velocity: **p=0.0013 for A53T vs A53T-Grm4 KO. **q**, Graph shows mean ± SEM of the latency to fall in the rotarod test in A53T-*Grm4*-/- mice. One-way ANOVA with Tukey’s post hoc test.

**Extended Data Figure 7.**
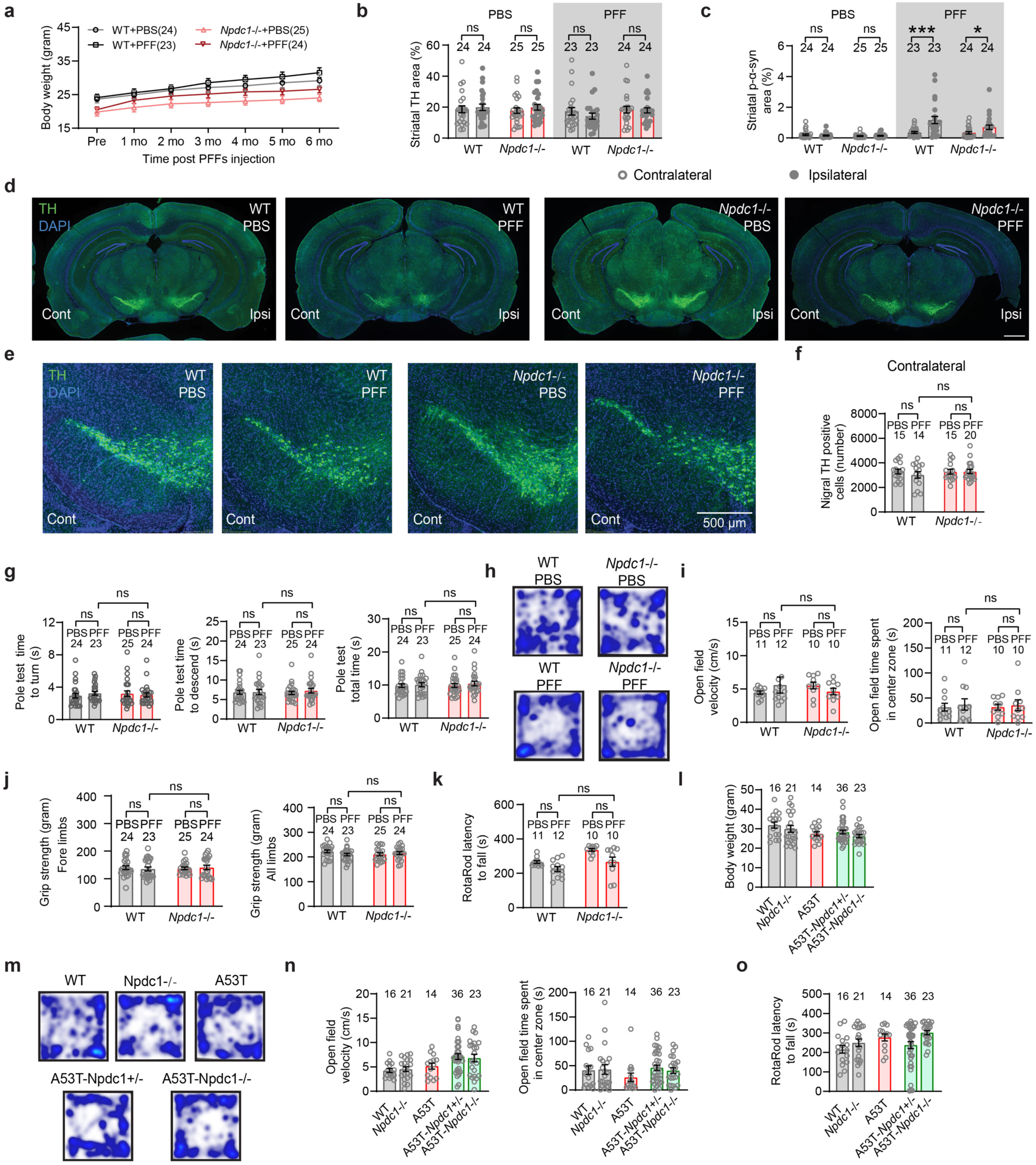
Additional phenotypic measures in *Npdc1*-/- mice injected with α-syn PFF and A53T-*Npdc1*-/- transgenic mice. **a,** Graph shows mean ± SEM of the body weight in *Npdc1*-/- mice injected with α-syn PFF. **b,** Graph shows mean ± SEM of the TH area in the ipsilateral and contralateral sides of the SNc in *Npdc1*-/- mice injected with α-syn PFF. Two-way ANOVA with Sidak’s post hoc test. **c,** Graph shows mean ± SEM of the p-α-syn area in the ipsilateral and contralateral sides of the SNc in *Npdc1*-/- mice injected with α-syn PFF. Two-way ANOVA with Sidak’s post hoc test. ***p<0.0001 for WT-PFF contralateral vs WT-PFF ipsilateral; *p=0.045 for Npdc1 KO-PFF contralateral vs Npdc1 KO-PFF ipsilateral. **d-e,** Representative images of TH (green) and DAPI (blue) staining in (**d**) the entire brain and (**e**) contralateral side of the SNc in *Npdc1*-/- mice injected with α-syn PFF. In **e**, coronal section with dorsal up and midline to right. Scale bars = 500 µm. **f,** Graph shows mean ± SEM of the number of TH positive neurons in *Grm4*-/- mice injected with α-syn PFF. One-way ANOVA with Tukey’s post hoc test. *p=0.018 for WT-PBS vs WT-PFF. **g,** Graphs show mean ± SEM of the time to turn (left), time to descend (center) and total (right) time in the pole test in *Npdc1*-/- mice injected with α-syn PFF. **h,** Representative images of the open field performance in *Npdc1*-/- mice injected with α-syn PFF. **i,** Graphs show mean ± SEM of the velocity (left) and the time spent in the center (right) in the open field test. One-way ANOVA with Tukey’s post hoc test. **j**, Graphs show mean ± SEM of the grip strength of the forelimbs (left) and all limbs (right) in *Npdc1*-/- mice injected with α-syn PFF. One-way ANOVA with Tukey’s post hoc test. **k**, Graph shows mean ± SEM of the latency to fall in the rotarod test in *Npdc1*-/- mice injected with α-syn PFF. One-way ANOVA with Tukey’s post hoc test. **l**, Graph shows mean ± SEM of the body weight in *Npdc1*-/- mice injected with α-syn PFF. One-way ANOVA with Tukey’s post hoc test. **m,** Representative images of the open field performance in A53T-*Npdc1*-/- mice. **n,** Graphs show mean ± SEM of the velocity (left) and the time spent in the center (right) in the open field test. One-way ANOVA with Tukey’s post hoc test. **o**, Graph shows mean ± SEM of the latency to fall in the rotarod test in A53T-*Npdc1*-/- mice. One-way ANOVA with Tukey’s post hoc test.

**Extended Data Figure 8.**
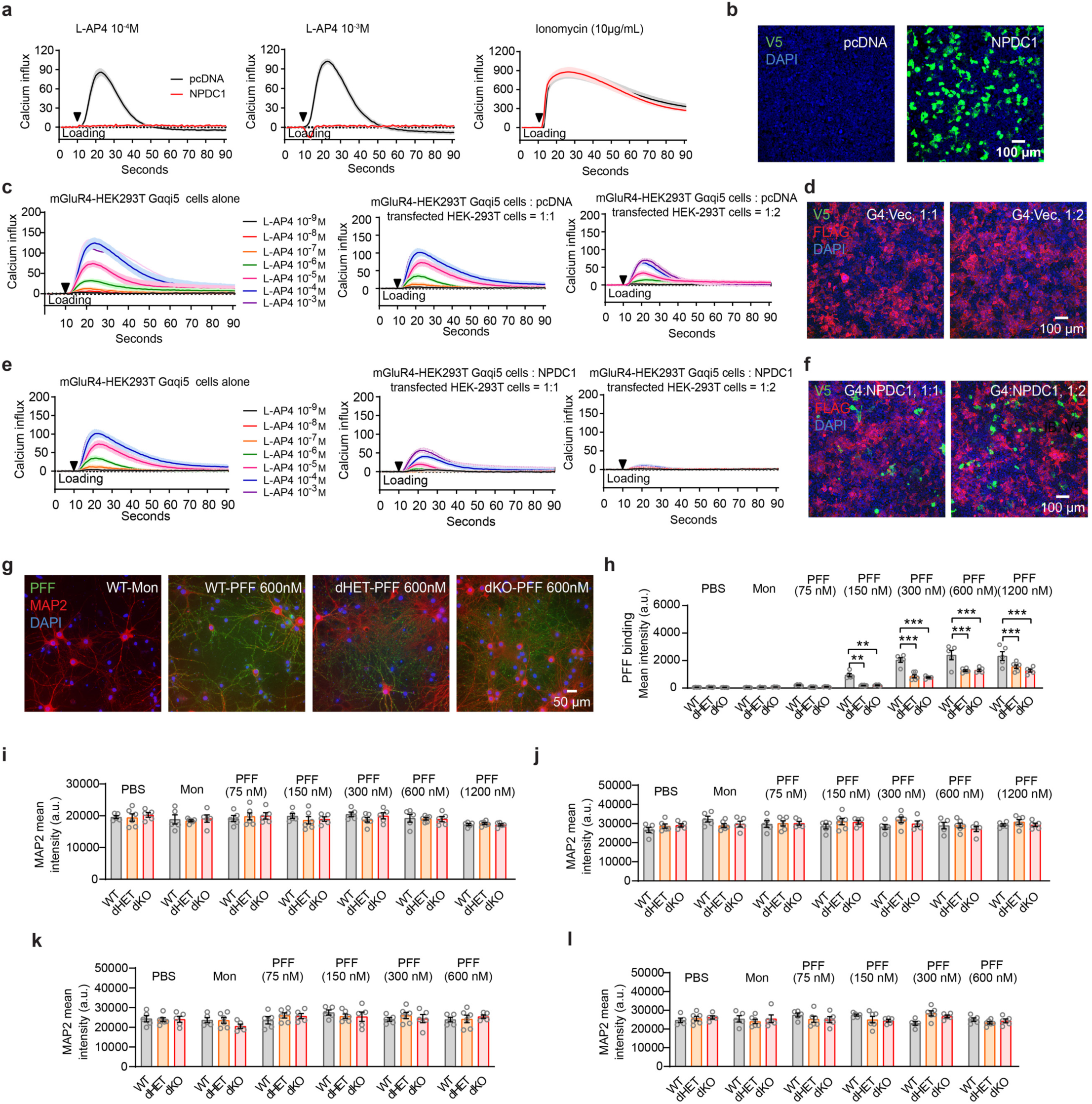
NPDC1 expression inhibits L-AP4 induced calcium influx in GRM4–Gαqi5 (G4)-HEK293T cells. **a,** Graphs show mean ± SEM of the calcium influx in GRM4–Gαqi5 cells transfected with an empty vector (pcDNA) or *NPDC1* and stimulated with 10^-4^ M (left) or 10^-3^ M (center) L-AP4 or ionomycin (right). **b,** Representative images of V5 (NPDC1, green) and DAPI (blue) staining in GRM4–Gαqi5 cells transfected with an empty vector (pcDNA) or *NPDC1.* Scale bar = 100 µm. **c,** Graphs show mean ± SEM of the calcium influx in GRM4–Gαqi5 cells alone (left) or co-cultured with HEK293T cells transfected with an empty vector (pcDNA) at a 1:1 ratio (center) and 1:2 ratio (right). Cells were stimulated with increasing concentrations of L-AP4. **d,** Representative images of V5 (NPDC1, green), FLAG (mGluR4, red) and DAPI (blue) staining in in GRM4– Gαqi5 cells co-cultured with HEK293T cells transfected with an empty vector (pcDNA) at a 1:1 ratio and 1:2 ratio. Scale bar = 100 µm. **e,** Graphs show mean ± SEM of the calcium influx in GRM4–Gαqi5 cells alone (left) or co-cultured with HEK293T cells transfected with *NPDC1* at a 1:1 ratio (center) and 1:2 ratio (right). Cells were stimulated with increasing concentrations of L-AP4. **f,** Representative images of V5 (NPDC1, green), FLAG (mGluR4, red) and DAPI (blue) staining in in GRM4– Gαqi5 cells co-cultured with HEK293T cells transfected with *NPDC1* at a 1:1 ratio and 1:2 ratio. Scale bar = 100 µm. **g,** Representative images of streptavidin (PFF, green), MAP2 (red) and DAPI (blue) staining of cultured neurons from WT, dHET and dKO mice after 2 hours of biotinylated α-syn monomer or PFF exposure at 4°C. Scale bar = 50 µm **h,** Graph shows mean ± SEM of the streptavidin (PFF) mean intensity in WT, dHET and dKO cultured neurons treated with different concentrations of α-syn PFF for 2 hours at 4 °C. One-way ANOVA with Tukey’s post hoc test. For 150 nM: **p=0.0011 for WT vs dHET, **p=0.0035 for WT vs dKO. For 300 nM: ***p<0.0001 for WT vs dHET, ***p<0.0001 for WT vs dKO. For 600 nM: ***p<0.0001 for WT vs dHET, ***p<0.0001 for WT vs dKO. For 1200 nM: ***p<0.0001 for WT vs dHET, ***p<0.0001 for WT vs dKO. **i, j,** Graphs show mean ± SEM of the MAP2 mean intensity in WT, dHET and dKO cultured neurons treated with different concentrations of α-syn PFF for 2 hours at 4 °C (i) or 37 °C (j). One-way ANOVA with Tukey’s post hoc test. **k, l,** Graphs show mean ± SEM of the MAP2 mean intensity in WT, dHET and dKO cultured neurons treated with different concentrations of α-syn PFF for 7 days at 4 °C (k) or 37 °C (l). One-way ANOVA with Tukey’s post hoc test.

**Extended Data Figure 9.**
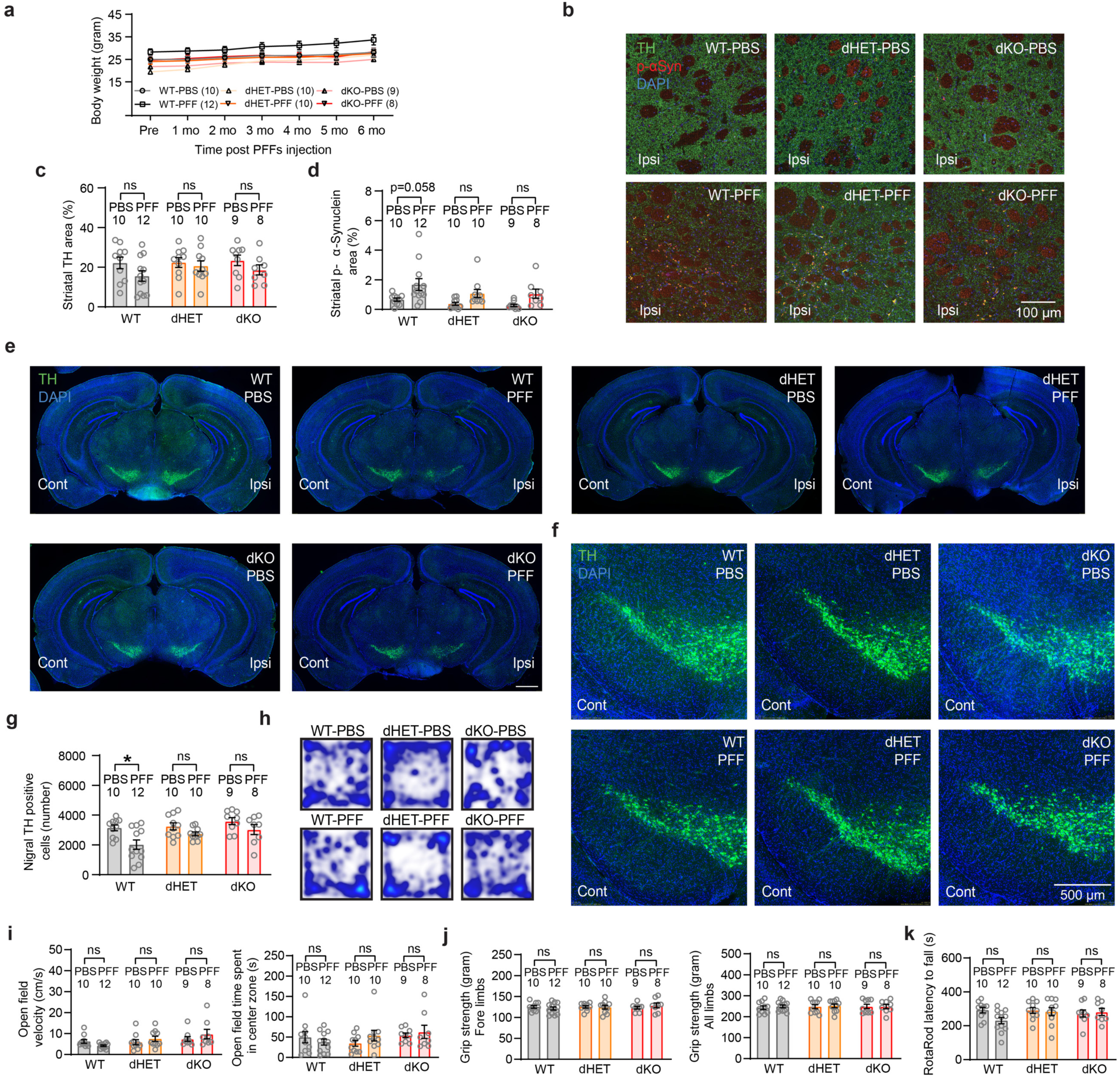
Additional phenotypes in Npdc1-Grm4 dHET and dKO mice injected with α-syn PFF. **a,** Graph shows mean ± SEM of the body weight in dHET and dKO mice injected with α-syn PFF. **b,** Representative images of TH (green), p-α-syn (red) and DAPI (blue) staining in the ipsilateral striatum of dHET and dKO mice injected with α-syn PFF. **c,** Graph shows mean ± SEM of the TH positive area in the ipsilateral striatum of dHET and dKO mice injected with α-syn PFF. One-way ANOVA with Tukey’s post hoc test. **d,** Graph shows mean ± SEM of the p-α-syn positive area in the ipsilateral striatum of dHET and dKO mice injected with α-syn PFF. One-way ANOVA with Tukey’s post hoc test. **e, f,** Representative images of TH (green) and DAPI (blue) staining in (**e**) the entire brain and (**f**) contralateral side of the SNc in dHET and dKO mice injected with α-syn PFF. In **e**, coronal section with dorsal up and midline to right. Scale bars = 500 µm. **g,** Graph shows mean ± SEM of the number of TH-positive neurons in contralateral SNc of WT, dHET and dKO mice injected with α-syn PFF. One-way ANOVA with Tukey’s post hoc test. *p=0.020 for WT-PBS vs WT-PFF. **h,** Representative images of the open field performance in dHET and dKO mice injected with α-syn PFF. **i,** Graphs show mean ± SEM of the velocity (left) and the time spent in the center (right) in the open field test in dHET and dKO mice injected with α-syn PFF. One-way ANOVA with Tukey’s post hoc test. **j**, Graphs show mean ± SEM of the grip strength of the forelimbs (left) and all limbs (right) in dHET and dKO mice injected with α-syn PFF. One-way ANOVA with Tukey’s post hoc test. **k**, Graph shows mean ± SEM of the latency to fall in the rotarod test in dHET and dKO mice injected with α-syn PFF. One-way ANOVA with Tukey’s post hoc test.

**Extended Data Figure 10.**
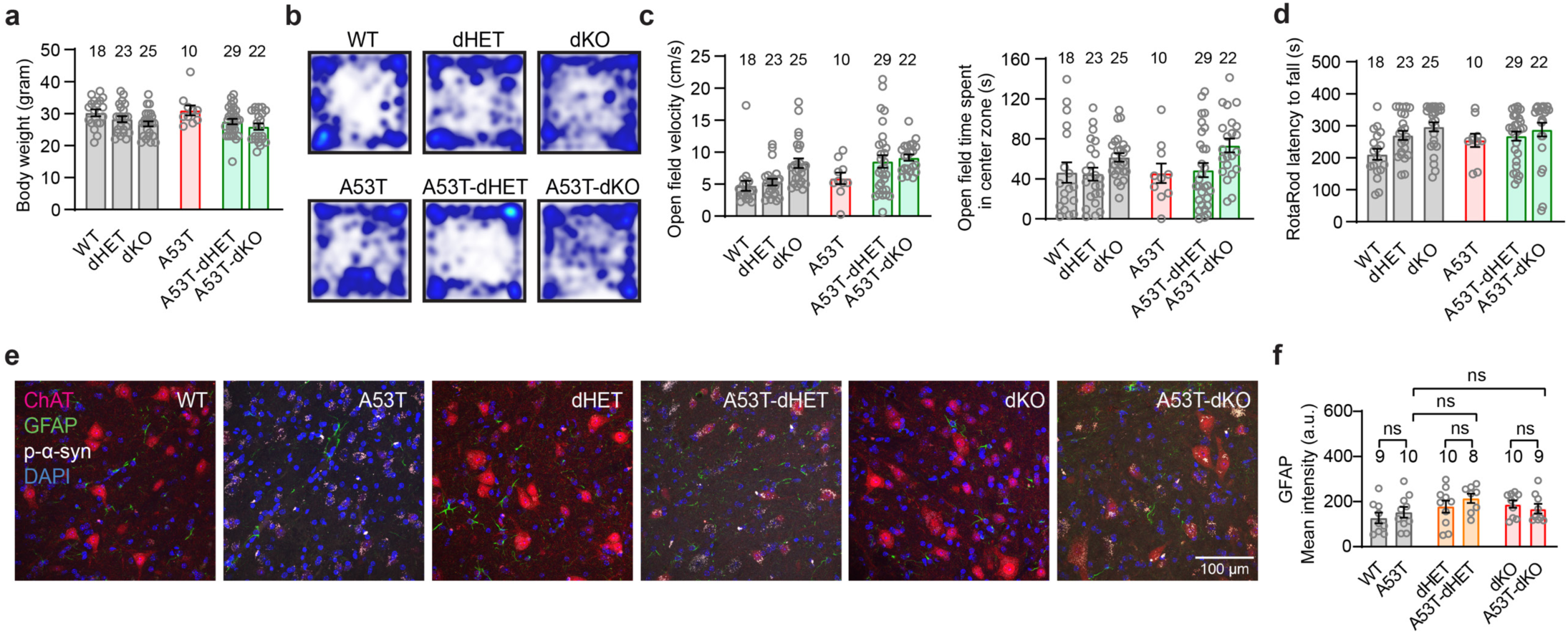
Additional phenotypes in A53T-dHET and A53T-dKO mice. **a,** Graph shows mean ± SEM of the body weight in A53T-dHET and A53T-dKO mice. **b,** Representative images of the open field performance in A53T-dHET and A53T-dKO mice. **c,** Graphs show mean ± SEM of the velocity (left) and the time spent in the center (right) in the open field test in A53T-dHET and A53T-dKO mice. One-way ANOVA with Tukey’s post hoc test. **d,** Graph shows mean ± SEM of the latency to fall from the rotarod test in A53T-dHET and A53T-dKO mice. One-way ANOVA with Tukey’s post hoc test. **e,** Representative images of ChAT (red), GFAP (green), p-α-syn (grey) and DAPI (blue) staining in the lumbar ventral horn of the spinal cord in A53T-dHET and A53T-dKO mice. Scale bar = 100 µm. **e,** Graph shows mean ± SEM of the GFAP mean intensity in A53T-dHET and A53T-dKO mice. One-way ANOVA with Tukey’s post hoc test test.

## Notes

### Competing Interest Statement

The authors have declared no competing interest.

## References

1. Spillantini, M.G., et al. Alpha-synuclein in Lewy bodies. Nature 388, 839–840 (1997).

2. Spillantini, M.G., Crowther, R.A., Jakes, R., Hasegawa, M. & Goedert, M. alpha-Synuclein in filamentous inclusions of Lewy bodies from Parkinson’s disease and dementia with lewy bodies. Proc Natl Acad Sci U S A 95, 6469–6473 (1998).

3. Dickson, D.W. Neuropathology of Parkinson disease. Parkinsonism Relat Disord 46 Suppl 1, S30–S33 (2018).

4. Goedert, M., Jakes, R. & Spillantini, M.G. The Synucleinopathies: Twenty Years On. J Parkinsons Dis 7, S51–S69 (2017).

5. Luk, K.C., et al. Pathological alpha-synuclein transmission initiates Parkinson-like neurodegeneration in nontransgenic mice. Science 338, 949–953 (2012).

6. Peng, C., et al. Cellular milieu imparts distinct pathological alpha-synuclein strains in alpha-synucleinopathies. Nature 557, 558–563 (2018).

7. Prusiner, S.B., et al. Evidence for alpha-synuclein prions causing multiple system atrophy in humans with parkinsonism. Proc Natl Acad Sci U S A 112, E5308–5317 (2015).

8. Masuda-Suzukake, M., et al. Prion-like spreading of pathological alpha-synuclein in brain. Brain 136, 1128–1138 (2013).

9. Neupane, S., De Cecco, E. & Aguzzi, A. The Hidden Cell-to-Cell Trail of alpha-Synuclein Aggregates. J Mol Biol 435, 167930 (2023).

10. Scheiblich, H., et al. Microglia rescue neurons from aggregate-induced neuronal dysfunction and death through tunneling nanotubes. Neuron 112, 3106–3125 e3108 (2024).

11. Henderson, M.X., et al. Spread of alpha-synuclein pathology through the brain connectome is modulated by selective vulnerability and predicted by network analysis. Nat Neurosci 22, 1248–1257 (2019).

12. Lauren, J., Gimbel, D.A., Nygaard, H.B., Gilbert, J.W. & Strittmatter, S.M. Cellular prion protein mediates impairment of synaptic plasticity by amyloid-beta oligomers. Nature 457, 1128–1132 (2009).

13. Mao, X., et al. Pathological alpha-synuclein transmission initiated by binding lymphocyte-activation gene 3. Science 353 (2016).

14. Gu, H., et al. Lymphocyte Activation Gene 3 (Lag3) Contributes to alpha-Synucleinopathy in alpha-Synuclein Transgenic Mice. Front Cell Neurosci 15, 656426 (2021).

15. Emmenegger, M., et al. LAG3 is not expressed in human and murine neurons and does not modulate alpha-synucleinopathies. EMBO Mol Med 13, e14745 (2021).

16. Ferreira, D.G., et al. alpha-synuclein interacts with PrP(C) to induce cognitive impairment through mGluR5 and NMDAR2B. Nat Neurosci 20, 1569–1579 (2017).

17. Thom, T., et al. Cellular Prion Protein Mediates alpha-Synuclein Uptake, Localization, and Toxicity In Vitro and In Vivo. Mov Disord 37, 39–51 (2022).

18. So, R.W.L., et al. alpha-Synuclein strain propagation is independent of cellular prion protein expression in a transgenic synucleinopathy mouse model. PLoS Pathog 20, e1012517 (2024).

19. Diaz-Ortiz, M.E., et al. GPNMB confers risk for Parkinson’s disease through interaction with alpha-synuclein. Science 377, eabk0637 (2022).

20. Brendza, R., et al. Genetic ablation of Gpnmb does not alter synuclein-related pathology. Neurobiol Dis 159, 105494 (2021).

21. Choi, Y.R., et al. FcgammaRIIB mediates the inhibitory effect of aggregated alpha-synuclein on microglial phagocytosis. Neurobiol Dis 83, 90–99 (2015).

22. Choi, Y.R., et al. Prion-like Propagation of alpha-Synuclein Is Regulated by the FcgammaRIIB-SHP-1/2 Signaling Pathway in Neurons. Cell Rep 22, 136–148 (2018).

23. Domingues, R., et al. Extracellular alpha-synuclein: Sensors, receptors, and responses. Neurobiol Dis 168, 105696 (2022).

24. Griesser, E., et al. Quantitative Profiling of the Human Substantia Nigra Proteome from Laser-capture Microdissected FFPE Tissue. Mol Cell Proteomics 19, 839–851 (2020).

25. Cenci, M.A., Skovgard, K. & Odin, P. Non-dopaminergic approaches to the treatment of motor complications in Parkinson’s disease. Neuropharmacology 210, 109027 (2022).

26. Rascol, O., et al. A Randomized, Double-Blind, Controlled Phase II Study of Foliglurax in Parkinson’s Disease. Mov Disord 37, 1088–1093 (2022).

27. Conn, P.J. & Pin, J.P. Pharmacology and functions of metabotropic glutamate receptors. Annu Rev Pharmacol Toxicol 37, 205–237 (1997).

28. Conklin, B.R., Farfel, Z., Lustig, K.D., Julius, D. & Bourne, H.R. Substitution of three amino acids switches receptor specificity of Gq alpha to that of Gi alpha. Nature 363, 274–276 (1993).

29. Galiana, E., Vernier, P., Dupont, E., Evrard, C. & Rouget, P. Identification of a neural-specific cDNA, NPDC-1, able to down-regulate cell proliferation and to suppress transformation. Proc Natl Acad Sci U S A 92, 1560–1564 (1995).

30. Evrard, C. & Rouget, P. Subcellular localization of neural-specific NPDC-1 protein. J Neurosci Res 79, 747–755 (2005).

31. Kim, S., et al. NGL family PSD-95-interacting adhesion molecules regulate excitatory synapse formation. Nat Neurosci 9, 1294–1301 (2006).

32. Pekhletski, R., et al. Impaired cerebellar synaptic plasticity and motor performance in mice lacking the mGluR4 subtype of metabotropic glutamate receptor. J Neurosci 16, 6364–6373 (1996).

33. Lee, M.K., et al. Human alpha-synuclein-harboring familial Parkinson’s disease-linked Ala-53 --> Thr mutation causes neurodegenerative disease with alpha-synuclein aggregation in transgenic mice. Proc Natl Acad Sci U S A 99, 8968–8973 (2002).

34. Martin, L.J., et al. Parkinson’s disease alpha-synuclein transgenic mice develop neuronal mitochondrial degeneration and cell death. J Neurosci 26, 41–50 (2006).

35. Evrard, C., Caron, S. & Rouget, P. Functional analysis of the NPDC-1 gene. Gene 343, 153–163 (2004).

36. Goedert, M. NEURODEGENERATION. Alzheimer’s and Parkinson’s diseases: The prion concept in relation to assembled Abeta, tau, and alpha-synuclein. Science 349, 1255555 (2015).

37. Moehle, M.S., et al. LRRK2 inhibition attenuates microglial inflammatory responses. J Neurosci 32, 1602–1611 (2012).

38. Doller, D., Bespalov, A., Miller, R., Pietraszek, M. & Kalinichev, M. A case study of foliglurax, the first clinical mGluR4 PAM for symptomatic treatment of Parkinson’s disease: translational gaps or a failing industry innovation model? Expert Opin Investig Drugs 29, 1323–1338 (2020).

39. Charvin, D., et al. An mGlu4-Positive Allosteric Modulator Alleviates Parkinsonism in Primates. Mov Disord 33, 1619–1631 (2018).

40. Marino, M.J., et al. Allosteric modulation of group III metabotropic glutamate receptor 4: a potential approach to Parkinson’s disease treatment. Proc Natl Acad Sci U S A 100, 13668–13673 (2003).

41. Beurrier, C., et al. Electrophysiological and behavioral evidence that modulation of metabotropic glutamate receptor 4 with a new agonist reverses experimental parkinsonism. FASEB J 23, 3619–3628 (2009).

42. Tomioka, N.H., et al. Elfn1 recruits presynaptic mGluR7 in trans and its loss results in seizures. Nat Commun 5, 4501 (2014).

43. Dunn, H.A., Patil, D.N., Cao, Y., Orlandi, C. & Martemyanov, K.A. Synaptic adhesion protein ELFN1 is a selective allosteric modulator of group III metabotropic glutamate receptors in trans. Proc Natl Acad Sci U S A 115, 5022–5027 (2018).

44. Volpicelli-Daley, L.A., Luk, K.C. & Lee, V.M. Addition of exogenous alpha-synuclein preformed fibrils to primary neuronal cultures to seed recruitment of endogenous alpha-synuclein to Lewy body and Lewy neurite-like aggregates. Nat Protoc 9, 2135–2146 (2014).

45. Hu, F., et al. Sortilin-mediated endocytosis determines levels of the frontotemporal dementia protein, progranulin. Neuron 68, 654–667 (2010).

